# Gaussian process forecasting of sparse ecological time series

**DOI:** 10.1101/2025.07.10.664121

**Authors:** Parul V. Patil, Robert B. Gramacy, Leah R. Johnson

**Affiliations:** Department of Statistics, Virginia Tech, Blacksburg, Virginia 24061, USA

**Keywords:** *Amblyomma americanum*, Ecological forecasting, heteroskedastic Gaussian process, Lone-star ticks, NEON forecasting challenge, Tick populations

## Abstract

Ecological time series are often unevenly sampled in time. That is, because the sampling processes used are resource intensive, data may be collected infrequently, or with adaptive frequencies triggered by presence of a target variable. When the data are irregularly spaced, standard time series methods may not be directly applicable. Instead, approaches that take inspiration from linear regression (LR) may be appropriate. In this paper, we explore flexible, nonparametric Gaussian process (GP) models as tools for producing forecasts of unevenly sampled observations. Our example is data on abundances of nymphal *Amblyomma americanum* from nine locations spread across the eastern United States, collected by NEON. The data exhibit highly variable sampling regimes and abundance levels across locations and time. We implement two versions of GPs to forecast tick abundance and benchmark models against LR approaches. Both GPs are able to capture population patterns without the need for forecasting additional drivers, such as temperature, or specifying a specific relationship between the response and predictors. We find that GP models provide an effective method to forecast irregularly sampled populations at short to intermediate time scales, outperforming LR and other comparators.

**Open Research statement:** This paper uses data that are already published and publicly available together with novel code. Both are provided together, to allow for reproduction of results, at a public git repository linked here: https://bitbucket.org/parulpatil22/tickgp-code/.

## 1 Introduction

Accurate forecasts of ecological time series are important for a variety of applications. For example, forecasts of algal blooms can ensure proper resource management as well as drinking water quality (Carey et al., 2022) and pest forecasting can allow farmers to spray crops with pesticides in a timely manner improving efficacy while reducing the risk of adverse health effects (Liu et al., 2022; Ponomarev et al., 2019). Forecasting of species dynamics over medium to long-term horizons can also be used to inform policy, for example, forecasting the effects of climate change on *Aedes*-borne viruses to inform public health measures (Ryan et al., 2019) or predicting the growth of chytrid fungus in frogs to help prevent rapid decline of amphibians (Gajewski et al., 2021). Although forecasting populations of species is useful in many domains, it can be challenging as monitoring and sampling populations can be difficult. For instance, observing endangered frogs affected by chytrid in the Sierra Nevada mountains of California requires researchers to haul sampling equipment on foot for hours to collect a single data point. As a result, populations may be irregularly and infrequently sampled which presents challenges for forecasting efforts.

One example of irregularly sampled populations for which short to medium term forecasts would be useful for making decisions are ticks that spread infectious diseases. In the United States and around the world, ticks adversely affect humans and animals, primarily by spreading tick-borne pathogens (Mitchell et al., 2020; Levin et al., 2017; Paddock et al., 2016; Piesman and Eisen, 2008). In the United States, diseases caused by tick-born pathogens include Lyme disease and Babesiosis, both transmitted by the blacklegged tick, *Ixodes scapularis*; and human Ehrlichiosis (Gaines et al., 2014; Raghavan et al., 2014; Childs and Paddock, 2003), Heartland virus (Savage et al., 2013), and Tularemia (Goddard and Varela-Stokes, 2009) (among others) transmitted by the *Amblyomma americanum* (commonly referred to as the lone-star tick). Apart from spreading pathogens, bites from ticks can also cause long term issues in humans such as alpha-gal syndrome which can result in a life long meat allergy (Steinke et al., 2015; van Nunen, 2015; Commins et al., 2011). Predicting tick density, as a proxy for bite risk, can help assess likelihood of pathogen transmission (Léger et al., 2013; Pfäffle et al., 2013). This can be beneficial for resource management, public safety, and preventative health measures (Estrada-Peña, 2001a; Randolph and Rogers, 1997)

To forecast tick density we rely on samples of observed tick activity. Ticks can be sampled in a variety of ways such as tick drags, CO_2_ traps, and baited traps. The trap methods are easier as they require less time and effort, but are not as effective as tick drags (Falco and Fish, 1992). The drag method involves an experimenter dragging a cloth across a study area to collect ticks. These are then identified to species or genus level and counts are reported. This method is time and effort intensive, and so adaptive sampling schedules are often used to minimize the chance that a drag is conducted when ticks are not present and active. This can lead to uneven sampling intervals and result in a small dataset with irregular gaps between sampling times. For example, our motivating dataset of tick observations from the National Ecological Observatory Network (NEON) has 385 observations across nine different locations over the past decade. Ideally, the number of observations (assuming weekly resolution) would be 53 weeks × 10 years × 9 locations ≈ 4,770 observations – an order of magnitude larger than the number of observations we have. This sparsity can make modeling challenging.

Classical time series (TS) models are a popular choice for forecasting (Brugger et al., 2018; Meltzer and Norval, 1992). TS models decompose observations into trend, seasonal, autoregressive (AR), and noise components. The AR term captures dependence between observations separated by a fixed lag which implicitly assumes regular sampling. In situations with sparse data, successive observations might be separated by variable time intervals. TS models for irregularly spaced data are an option for fitting irregular time seris (Robinson, 1977), however the estimation of an AR term requires extensive data pre-processing including imputation and aggregation over large time intervals. This can dampen the signal and lead to biased estimates of AR coefficients (Little and Rubin, 2019). Identifying seasonality parameters can also be challenging as the phase is not always identifiable due to large gaps. In such situations, classical time series models that rely on evenly spaced data may result in misleading forecasts for unevenly sampled data (Johnson and Munch, 2022).

State space models (SSMs) are also commonly used for modeling irregularly sampled time series (Newman et al., 2023; Auger-Méthé et al., 2021; Jones, 1986). These models are hierarchical and explicitly separate the unobserved latent process (which models the underlying dynamics) from the observed process that is subject to measurement error (Cressie et al., 2009). SSMs depend on a resolution of observations relative to the resolution at which we want to infer the true process (Auger-Méthé et al., 2021). When there are large gaps in the data, the observations may not provide enough information to infer latent states. For example, changes in observations could be due to several factors such as fast-changing dynamics, high process noise, or measurement errors which the model cannot distinguish amongst without prior information (Auger-Méthé et al., 2016). Since parameter estimation is challenging, inference relies on predicting latent states forward in time rather than updating it through the data. Errors in estimation propagate through the model and can produce unreliable estimates that depend strongly on the prior.

We instead propose a Gaussian process (GP) model, which is a non-parametric regression model that leverages relative distances in the input space to inform predictions (Gramacy, 2020; Cressie, 1991). This approach incorporates a noise parameter to separate the mean process (here tick abundance) from the noise (i.e., sampling error, labeling error, etc). The standard formulation of GPs is homoskedastic, that is, the noise process is assumed to be the same in all parts of the input space. However, for our application, we expect that one estimate of noise across all locations would be insufficient and we instead prefer a heteroskedastic GP (hetGP) model. This variant allows both the mean and noise process to vary across the input space. Our contribution is not to simply say use a GP/hetGP and make predictions for tick applications. Instead we propose a framework that checks the following boxes: (1) It relaxes the assumption that the time series must be evenly sampled and is applicable to very sparse datasets such as ours. (2) It does not use environmental/weather predictors (e.g., temperature, humidity) that themselves need to be forecast, nor assume specific functional forms between the response and predictors. (3) It trains one hierarchical model across all locations to allow information sharing rather than training and forecasting individually at each location, as some locations are very sparse compared to others.

The main factor which determines performance of our GP models are the predictors that we choose. For a GP to make meaningful forecasts, we need well defined “closeness” in our input space. Models which determine presence/absence of ticks can provide informative features/predictors for our use. For example, classification models such as generalized linear models (GLM), maximum entropy models (MaxEnt; Raghavan et al., 2016), and classification and regression trees (CART – which are usually used in the field of machine learning (Zannou et al., 2021)) often incorporate weather and climate data such as temperature, moisture, and precipitation (Williams et al., 2015; Springer et al., 2015; Estrada-Peña, 2001b; Haile and Mount, 1987). Additionally, they can incorporate location-based features such as elevation and forest cover (Slatculescu et al., 2020; Hahn et al., 2016) and host density (Wimberly et al., 2008) which may influence tick abundance (Paddock and Yabsley, 2007). We determine some of our predictors based on features used in these other models. Note that position of features in the input space itself does not directly affect GP performance. Rather, the distance between them is what maters. An absolute value of temperature (or forecasts) are not required, only general patterns. We detail our predictors later in Sec. 3.2.

To benchmark our method, we train additional models as comparators to our GP models. In order to avoid the need to aggregate or interpolate, we do not build TS models and instead focus on two types of comparators. First are a set of linear regression (LR) models with simple predictors. Second are Bayesian adaptive spline surfaces (BASS) which is a common flexible and non-parametric regression framework designed for nonlinear relationships (Francom and Sansó, 2020; DiMatteo et al., 2001). BASS models directly attempt to infer the underlying signal given the data without assuming a time-series structure and adjust the smoothing based on the data (Serra and Krivobokova, 2017; Wood et al., 2002). They can thus model sparse data without requiring imputation. We do not describe BASS in detail (see cites for more) and simply use it as a comparator to our GP methods. To level the playing field, GPs and BASS use the same predictors.

Below, we begin by introducing the problem setup with details of the data source followed by visualization and pre-processing steps (Section 2). Section 3 covers various modeling approaches starting with a linear regression (LR) comparator, which representing a classic approach to this problem (Section 3.1). We then introduce the GP model along with implementation details (Section 3.2) and an extension, namely HetGPs which is our proposed approach for this specific forecasting problem (Section 3.3). Section 4 consists of comparisons of all the modeling approaches across multiple metrics. Lastly, we conclude with a brief discussion of the methods and the future scope (Section 5).

## 2 Data Description and Preparation

### 2.1 Data Overview

#### 2.1.1 Amblyomma americanum

In this study, we focus on modeling the seasonal density of *Amblyomma americanum. Amblyomma* ticks follow a three-host life cycle (larval, nymphal, and adult stages – each on a different animal host) that spans approximately 8 months to 2 years, depending on environmental conditions (Troughton and Levin, 2007). At each stage, the tick requires a blood meal from a host, typically mammals, birds, rodents, or humans, to develop further (Kollars Jr. et al., 2000). Feeding usually lasts one to three weeks per stage (Troughton and Levin, 2007). *A. americanum* ticks specifically inhabit wooded areas and primarily feed on white-tailed deer (Allan et al., 2010; Paddock and Yabsley, 2007), though wild turkeys are common hosts for immature stages, especially in the Midwest (Mock et al., 2001). They also aggressively quest for hosts based on carbon-dioxide and movement patterns (Childs and Paddock, 2003).

*A. americanum* ticks are native to the eastern and southeastern United States and parts of Mexico (Molaei et al., 2019; Monzón et al., 2016; Springer et al., 2014). However, their range has been expanding due to climate change (Rochlin et al., 2022; Raghavan et al., 2019; Cortinas and Spomer, 2013; Masters et al., 2008). Notably, there have been reports of these ticks in northern regions of the U.S. that were historically too cold for their survival (Paddock and Yabsley, 2007). This expansion is concerning, as *A. americanum* are responsible for a significant proportion of tickborne diseases in North America (Goddard and Varela-Stokes, 2009; Mixson et al., 2006; Goddard, 2002; Felz et al., 1996).

#### 2.1.2 NEON Forecasting Challenge

Our forecasting effort was based on a subset of tick data obtained from National Ecological and Observatory Network (NEON) aggregated for a tick forecasting challenge set up by Ecological Forecasting Initiative Research Coordination Network (EFI-RCN; Thomas et al., 2023). The target data are counts of nymphal *A. americanum* ticks and also includes the date of collection, the location, the target species, and the iso-week of collection. As part of the NEON challenge, we also have a subsidiary dataset (site dataset; Site Metadata, 2025), with information about the nine locations such as the coordinates, altitude (mean, minimum and maximum), and ecological features such as precipitation, temperature, foliage, etc. In Fig. 1, we show the nine NEON locations along with mean tick density observed at each location in the past decade and total number of observations at each location.

**Figure 1.**
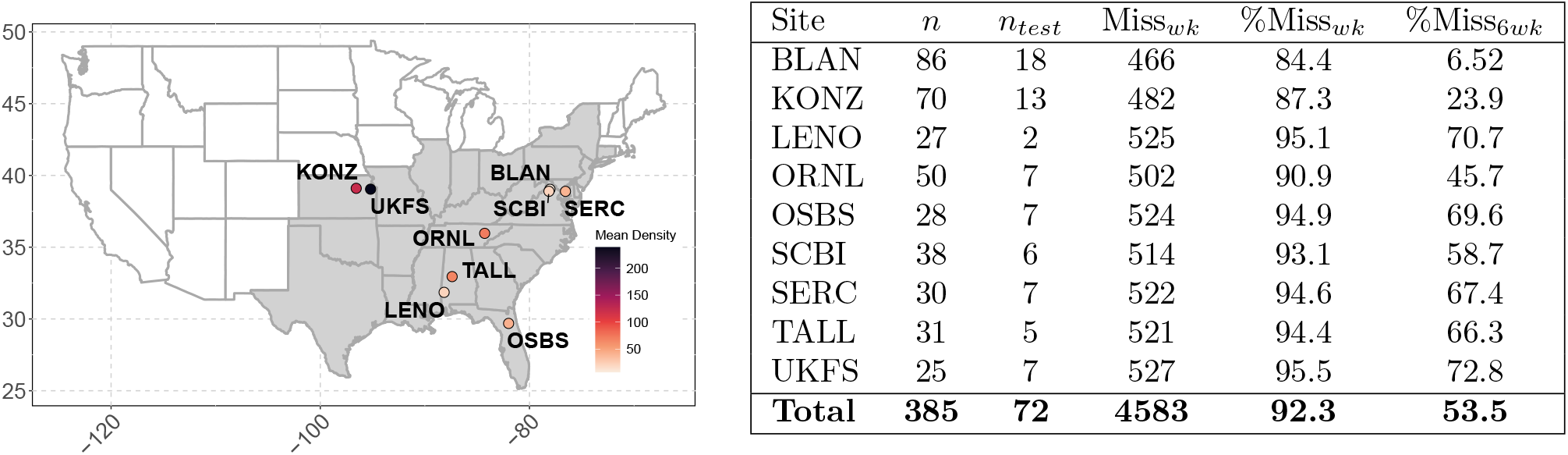
*(left)* NEON locations with average tick density at different locations. *(right)* Details such as number of samples collected, used for testing, along with number and % of missing observations.

Ticks are sampled at NEON locations using the “tick drag” method. Specifically a 1*m*^2^ cloth is manually dragged across the plots in transects of 160m (National Ecological Observatory Network (NEON), 2024). This occurs once every few weeks at a given location during the sampling season which is determined by the foliage at the location. The collected ticks are then taken to the lab for taxonomic identification where the life stage and the taxonomic rank of the species is identified. In Fig. 1 (*right)*, we have provided observed and missing samples for each location assuming a weekly sampling scheme. Notice that almost 90% of data is missing. Assuming a highly adaptive sampling scheme every six weeks, we would still be missing about 50 % data at all locations. This can present as a challenge especially considering that forecasts are required at weekly resolution.

#### 2.1.3 Data Visualization

In this section, we highlight key exploratory observations across locations that inform our modeling decisions. We illustrate the location-level variation in Fig. 2 by showing tick density recorded at three representative sites: Lenoir Landing (LENO), Smithsonian Conservation Biology Institute (SCBI), and Talladega National Forest (TALL). Tick samples vary across locations due to difference in sampling schemes. For example, samples for some locations, e.g., TALL/SCBI date back to 2014, while LENO only has data 2016 onwards. The sampling season also changes depending on geographic location. Some regions experience lower temperatures and more snowfall which hinders longer tick dragging schemes in winter. However, warmer regions (SCBI/BLAN) do have occasional drags in winter as ticks can remain active year round.

**Figure 2.**
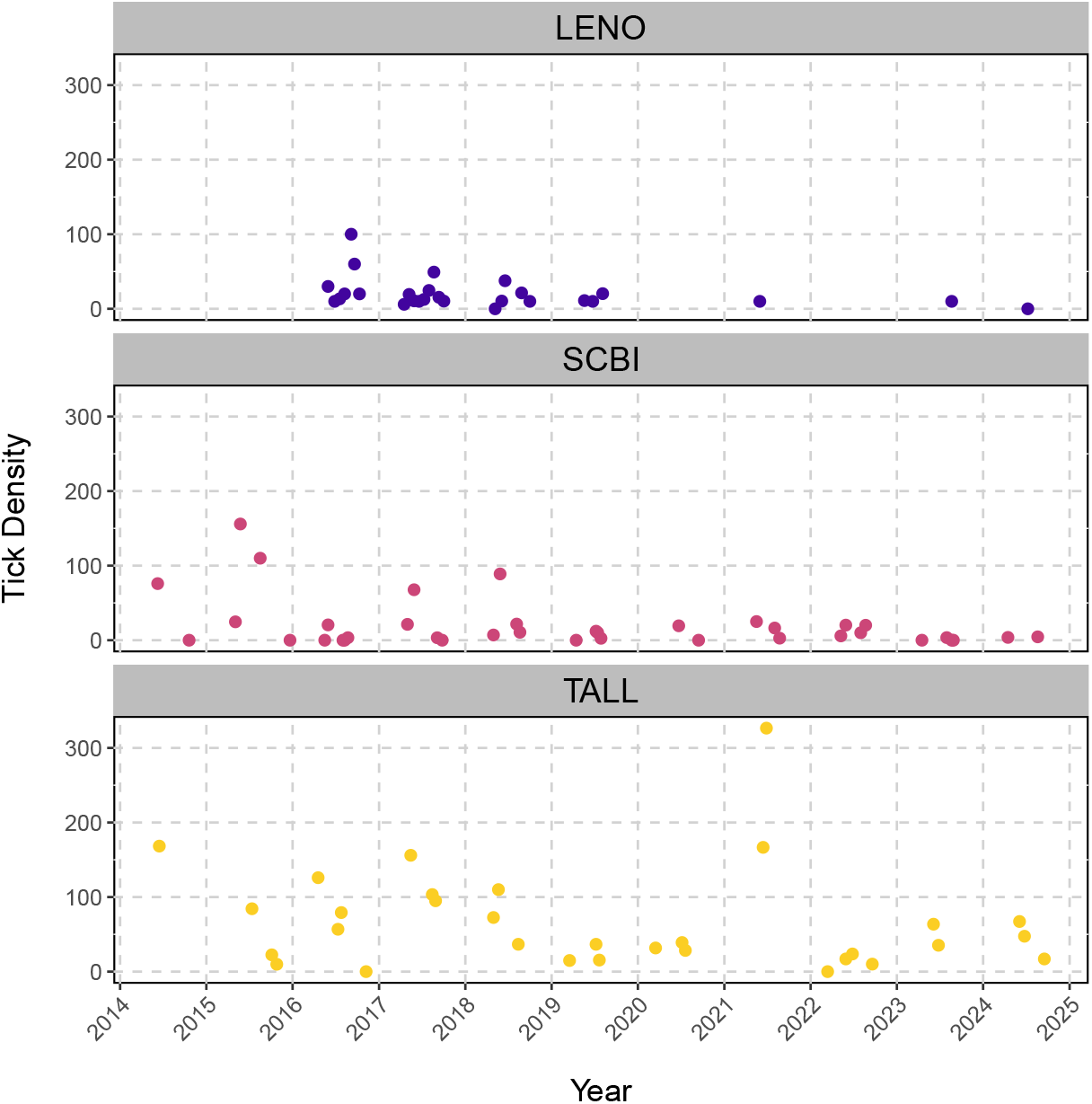
Observed Tick Density across three different NEON locations in the past decade

We also observe a difference in the amplitude of tick density across locations. For instance, peak densities at TALL reach around 300 ticks per plot, while LENO peaks at approximately 100 ticks per plot. Other sites observe over 600 ticks per plot during peak season. We can also identify some seasonal patterns in Fig. 2. Observed density is lower in colder months and higher during warmer months at each location. However, the magnitude of seasonal peaks can vary significantly from year to year, even at the same site.

### 2.2 Preprocessing Steps

#### 2.2.1 Data Transformation

To forecast tick density, we explore modeling methods that rely on the assumption that the response must be normally distributed around the mean. Since tick density is strictly positive and includes true zeros, we use a transform to ensure this constraint is satisfied. We use a combination of the square-root and logarithm transform given by:

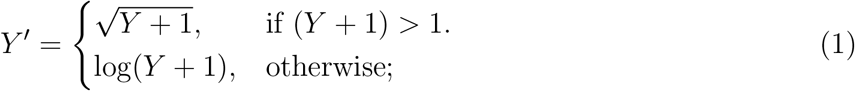

where, *Y* ^*′*^ is the transformed response and *Y* is on the original scale, i.e., the observed tick density. Notice that taking a log of zero is avoided by shifting tick density (add one to original density). Using a square-root transform for larger values prevents over expansion on the back transform (which would be a consequence of using only a log transform). Similarly, using a log transform for smaller values prevents insensible UQ (upper bound *<* lower bound) on the back transform.

#### 2.2.2 Train-Test Split

To evaluate our methods, we split the data into two sets, training and holdout (testing). For time series data, it is standard to use a cutoff date before which all the data is historic training data. Data after the cutoff is used as the holdout set to evaluate predictions. We selected a cutoff date of 2022-12-31 which provided a sufficient number of points to assess model performance. Because predictions made only at sparse time points do not capture underlying periodic trends well, we constructed a uniform weekly grid from 2014 to the latest date in the dataset. Each of the trained models was used to predict tick density at every week in this grid. Predictions corresponding to all the holdout dates were picked out to evaluate out-of-sample performance.

We benchmarked our method using coverage, root mean square error (RMSE; Appendix S1: Section A), and continuous rank probability score (CRPS; Gneiting and Raftery, 2007). Coverage is the percentage of data points within the 90% predictive intervals (PIs), which ideally should match the nominal level. RMSE measures prediction accuracy ignoring uncertainty, while CRPS accounts for both accuracy and uncertainty and can be computed in closed form for Gaussian distributions. We calculated RMSE and CRPS on the transformed scale for all models. Since we are working with a sparse dataset, RMSE and CRPS are based off very small sample sizes and may not sufficiently capture the model performance. For example, we expect ticks to be inactive in winter and peak in the summer, but some models might predict constant densities with large uncertainty through out the year and still score well on metrics at sparse testing locations. Thus, we also visually assess all the models. In the results below, we preferred to present selected model fits that best illustrate specific characteristics of each modeling approach (see Appendix S1: Section C for all model fits at all locations). Additionally, we induced some dummy reference points to better quantify accuracy. Specifically, we added a zero tick density observation in the middle of the non-sampling season which was typically the last week of January (for all years) at all locations. The addition of the reference points penalizes models which do not predict a zero density in winter and allows us to better quantify model fits.

## 3 Modeling

### 3.1 Linear regression modeling

We start with using linear regression (LR) since it offers a simple alternative to time series modeling when data are irregularly sampled. The models can be expressed as *Y* ^*′*^ = *Xβ* + *ϵ* where 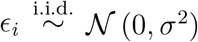. Here, *Y* ^*′*^ is the vector of responses (transformed tick density) of size *n* which corresponds to number of samples. The matrix *X* has dimensions *n ×* (*p* + 1), with *p* predictors appended with one column (of ones) corresponding to the model intercept. The parameter vector *β* = (*β*_0_, *β*_1_, …*β*_*p*_) includes the model intercept and *p* slopes. Random errors *ϵ*_*i*_ are assumed to be independent and identically distributed (i.i.d.), following a normal distribution with mean zero and constant variance *σ*^2^.

We discuss two variants of LR models distinguished by the primary predictor used in each case. The first model is the *LR-Time(L)* model which is our vanilla comparator. We use (L) to represent that the model was trained using a subset of data i.e., data which was collected at that specific **L**ocation rather than data collected at **A**ll locations (denoted by (A) in later sections). This model uses iso-week as a predictor with additional periodicity through sine and cosine predictors – each having a period of 53, representing the number of iso-weeks per year i.e., season in the context of our problem. Observe in Fig. 3a that the sine wave completes two cycles but we suspect that it is not in phase with tick density. Using both sine and cosine predictors allows the model to adjust the phasing. For our second variant, *LR-Temp(L)*, we used minimum temperature as a predictor rather than iso-week for reasons we explain shortly. All linear models are fit using the `stats` package in R (R Core Team, 2022).

**Figure 3.**
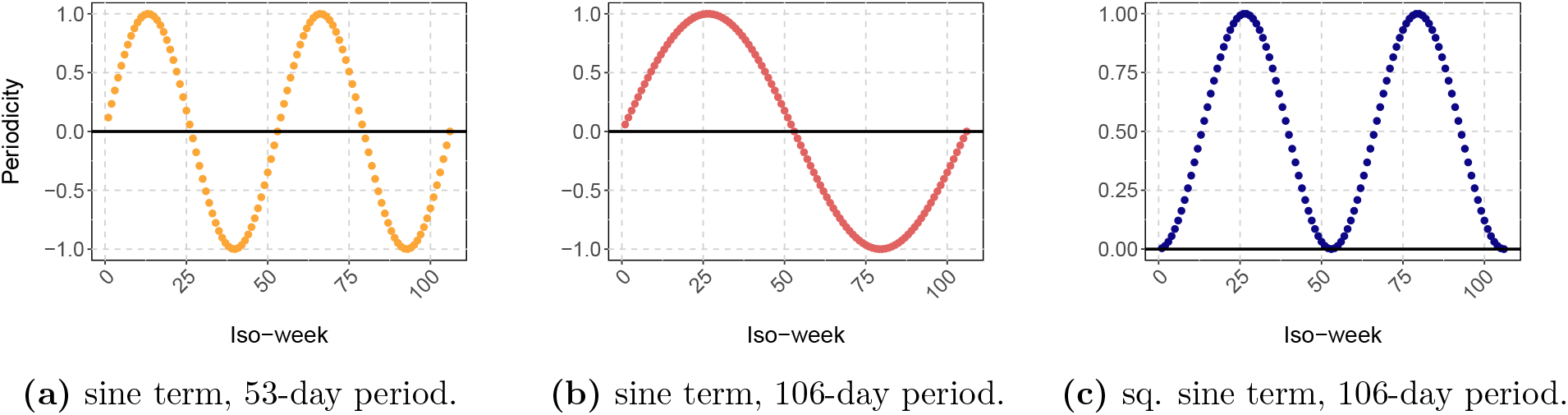
Periodicity for sine predictor with different phases and formulations explained above. We use (a) for linear models and (c) for Gaussian processes.

The predictors in the *LR-Time(L)* model repeat across years so we visualize the mean predictions as a function of iso-week. At both locations, i.e., UKFS and OSBS in Fig. 4, there is a discontinuity in the predictions across years and overly conservative upper bounds (Appendix S1: Section C). At UKFS (18 training points), the model captures the expected pattern of increase in ticks in summer and decrease in winter. However, at OSBS (21 training points), the model fails to capture this pattern, which we suspect is because the mode of tick-density changes from one season to another, making it challenging to identify the real trend across seasons using only iso-week and periodicity as predictors.

**Figure 4.**
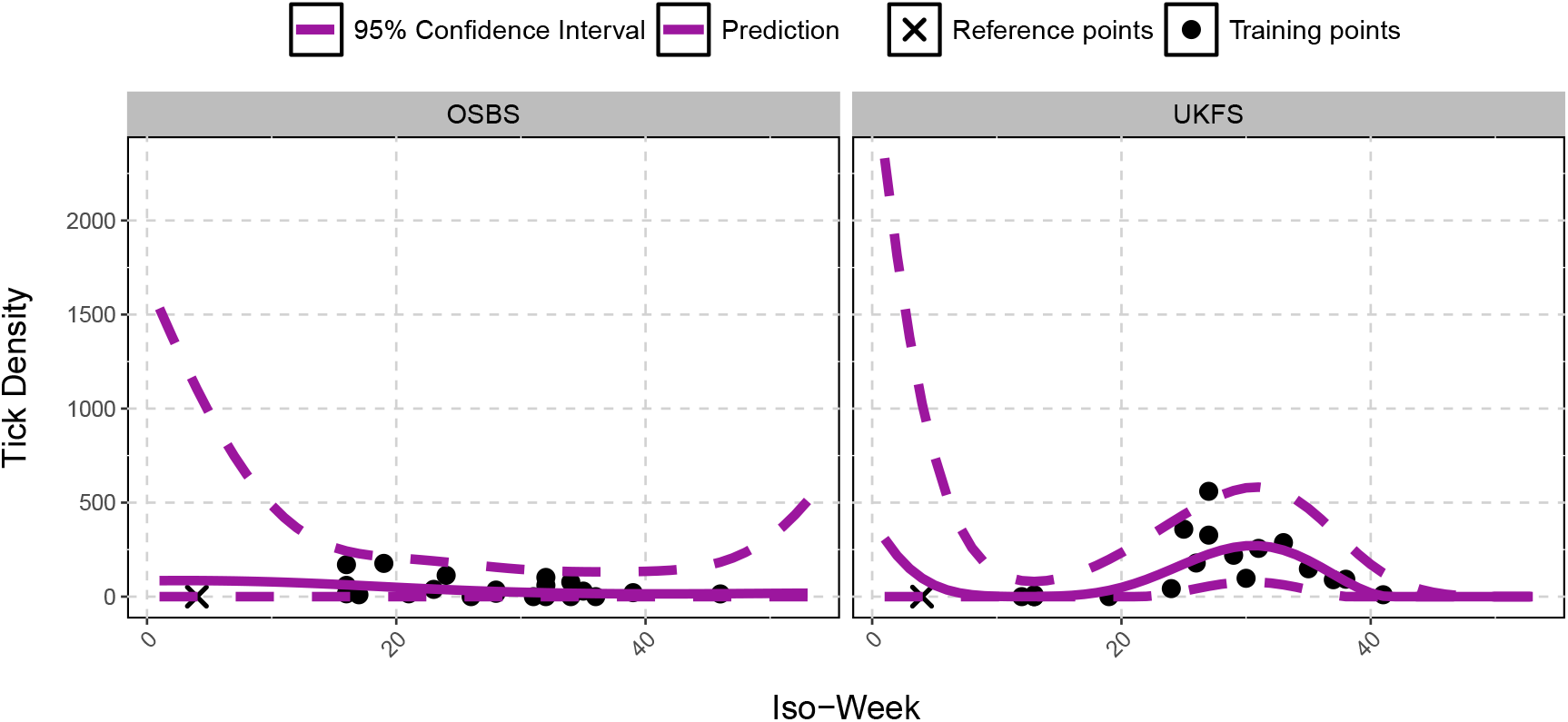
*LR-Time(L)* fit across iso-weeks 1–53 at (a) UKFS and (b) OSBS (solid = mean, dashed = 90% prediction intervals) on the original (untransformed) scale. Black dots show training data.

We now move onto our next variant which we call *LR-Temp(L)* model. We consider minimum temperature (*tmin*) as our predictor in place of iso-week for this model. In terms of modeling choices, we had a few options available to us. The first major choice was between weather data provided by NOAA Global Ensemble Forecasting System (Hamill et al., 2022)^1^ or historic monthly climatology data obtained from CRU-TS 4.09 (Harris et al., 2020) downscaled with WorldClim 2.1 (Fick and Hijmans, 2017). The main difference is that weather is often unstable over short periods and using weather forecasts for tick density estimation can lead to severely conservative uncertainty. Instead, we chose to work with climatology, averaging over short-term atmospheric conditions, providing less noisy, more stable forecasts.

The next choice was between different meteorological variables. We had data available for temperature, humidity, and precipitation, all of which are known to affect tick density (Clarke-Crespo et al., 2020; Sagurova et al., 2019; Raghavan et al., 2016; Vail and Smith, 1998). Although humidity and precipitation are important (Clarke-Crespo et al., 2020; Raghavan et al., 2016), they are difficult to measure and are highly correlated with temperature. Thus, we decided to use the minimum temperature as the predictor. We used climatology data from the years 2014 - 2024 at 5 mins resolution to obtain *tmin* (in Celsius) recorded at monthly frequency. To match the weekly frequency of our dataset, we repeat the same temperature for all weeks in the same month. In addition to this predictor, we also use the sine and cosine predictors as in the *LR-Time(L)* model.

We show an example fit of the *LR-Temp(L)* model obtained at location ORNL in Fig. 5. This model performs well on the training data where it predicts low tick density during colder months and captures uncertainty adequately. However, for true forecasts into the future, temperature is unknown and must be forecasted. Forecasting climate variables such as monthly average temperatures even one year in advance is inherently challenging due to short-term variability.

**Figure 5.**
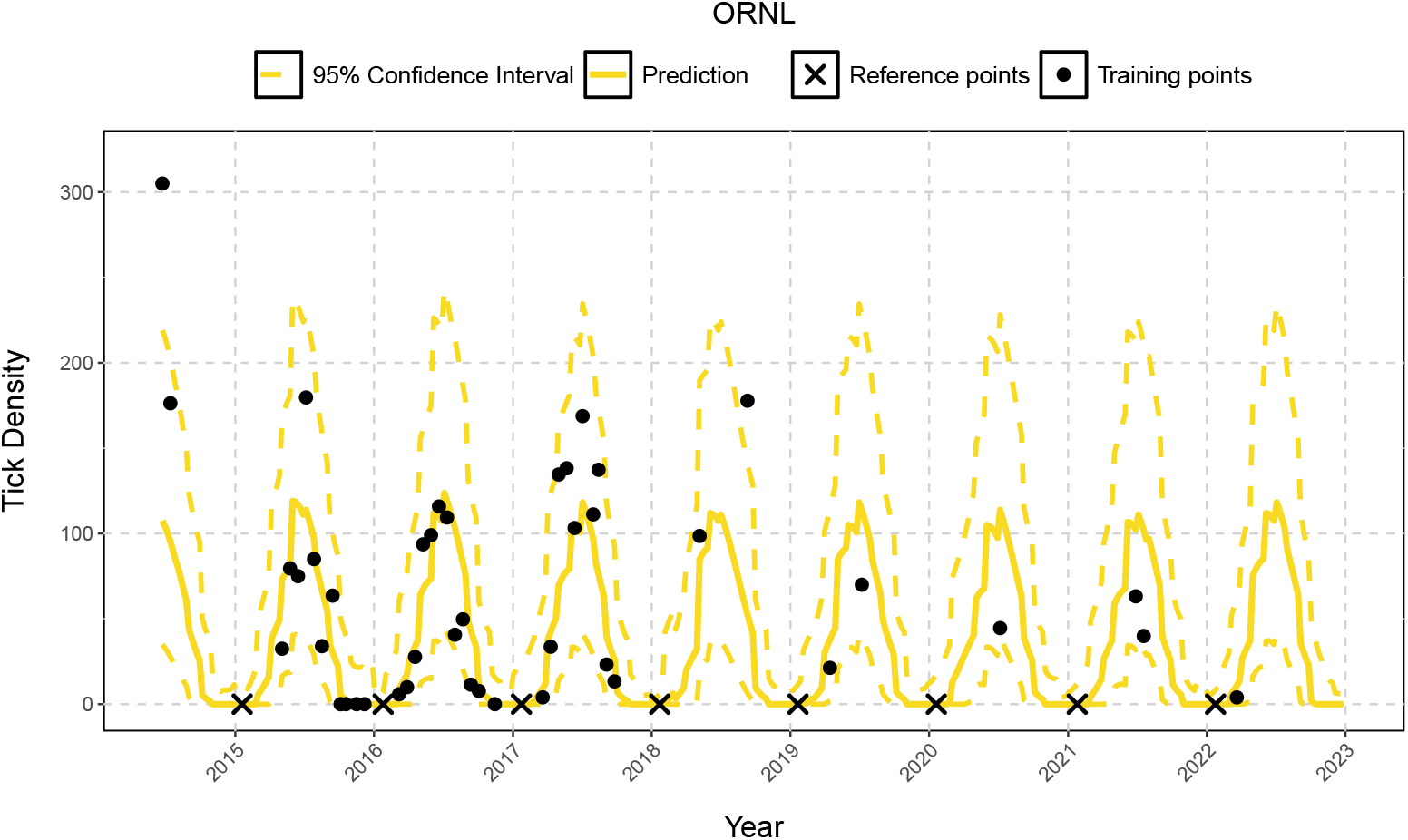
*LR-Temp(L)* fit at ORNL (solid = mean, dashed = 90% prediction intervals) on the original (untrasformed) scale. Black dots show training data.

### 3.2 Gaussian processes

In the previous sections, the LR models have served as a baseline for our analysis and provided insights into potential improvements. These include eliminating environmental factors, training one model for all locations, and obtaining better uncertainty quantification (UQ). In this section, we explore Gaussian processes (GPs; Gramacy, 2020) which are highly flexible and non-parametric models that can be used to learn patterns within data. GP regression models rely primarily on the distance between input values rather than their position, which has made them widely used in spatial statistics (where they are known as kriging; Cressie, 1991). GPs have also gained popularity in machine learning due to their ability to provide fast predictions at new locations along with reliable UQ (Rasmussen and Williams, 2005). We apply GPs in our ecological context because they do not require equally spaced time series, are flexible, and are particularly effective at distinguishing signal from noise (Johnson et al., 2018).

GP models are formulated as *Y* ∼ 𝒩 (*µ*_*n*_, Σ_*n*_). LR is technically a specific case of a GP where *µ*_*n*_ = (1, *X*_*n*_)*β* and Σ_*n*_ = *σ*^2^𝕀. However, in case of a GP, we focus specifically on the covariance structure Σ_*n*_ which is specified through a kernel. This kernel is built so that if training points *x*_*i*_ and *x*_*j*_ are closer together in the input space, the GP assumes responses *Y*_*i*_ and *Y*_*j*_ are more highly correlated. There are many kernels that fit this description, but in our case we use the squared exponential kernel *C*_*θ*_(*X*_*n*_) stated as

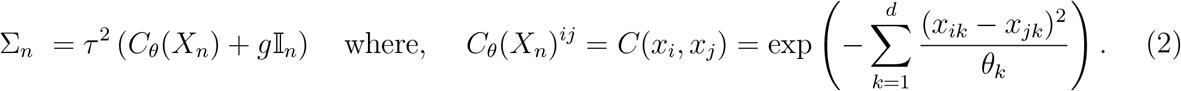

Through the kernel, the covariance structure is a function of three hyper-parameters: scale, *τ* ^2^ which captures the amplitude of the response; the length-scales, ***θ*** ={ *θ*_1_, *θ*_2_, …, *θ*_*d*_ } which captures the rate of decay of correlation for each dimension; and the nugget, *g*, which captures the noise observed in the response. Note that Eq. 2 represents an anisotropic specification where a unique length-scale is estimated for each dimension (as opposed to isotropic kernel where only one common length-scale is inferred for all dimensions.) We specify *µ*_*n*_ = 0 i.e., zero mean (for simplicity) and infer hyper-parameters {*τ* ^2^, ***θ***, *g*} through Maximum Likelihood Estimation (MLE).

We can then obtain the predictive distribution of 𝒴 at *n*_*p*_ new locations 𝒳 by calculating the conditional distribution of 𝒴, given (*X*_*n*_, *Y*_*n*_). To do so, first stack the distribution of 𝒴 and *Y*_*n*_ and jointly write it as,

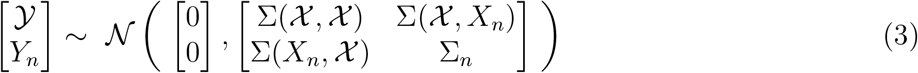

where 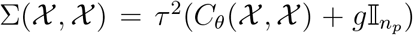 is the variance matrix for 𝒳 and Σ(*X*_*n*_, 𝒳 ) is the covariance matrix between 𝒳 and *X*_*n*_. Then, using MVN conditioning properties, we have 𝒴|*Y*_*n*_, *X*_*n*_ ∼ *N* (*µ*(𝒳 ), *σ*^2^(𝒳 )), and

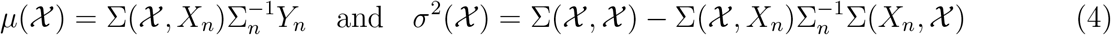

where *µ*(𝒳 ) is the mean and *σ*^2^(𝒳 ) is the variance at the predictive locations.

The bulk of our work is in building informative predictors *X*_*n*_ that capture “closeness” in our system. To build predictors, we have some criteria learnt from our previous sections, such as building predictors which do not require forecasting. Further, predictors should be able to capture temporal patterns along with geographical patterns, so we can train the GP model on a unified dataset, i.e., using data from all locations. We demonstrate shortly why this is crucial to obtain good fits in the context of sparse data.

Our first *common predictor* (the same for all locations) includes week number, 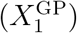 which is a grid of time points {1, 2, …, *n, n* + 1, …, *n* + *n*_*p*_} from the start date (week 1) up to the end date (week *n* + *n*_*p*_) where week numbers *n* and *n*_*p*_ correspond to the last week in the training and testing set respectively. By using week number as a predictor, we curate a relationship between the previous week and the current week since the distance between consecutive weeks will be the small. Next, we incorporate information that weeks across years should be similar through periodicity, 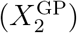. This predictor is based on the LR sine predictor, 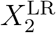, with some modifications i.e., increasing the frequency from 53 to 106 weeks (Fig. 3b) and squaring the sinusoid so we have 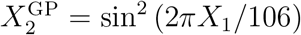. With these adjustment, the curve completes half a cycle in one year and flattens out towards the end of the year allowing for smoother transition between years.

In contrast to our common predictors above, *location specific predictors* are designed to distinguish one location from another. We obtained one location-specific predictor directly from the information provided by NEON namely, mean elevation 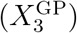 of location which is correlated with maximum tick density (Fig. 6a). The intuition behind this is that if observations *Y* and *Y* ^*′*^ from our unified dataset are collected at the different geographical locations but having similar elevation, the Euclidean distances between them would be small resulting in higher correlation through Σ_*n*_.

**Figure 6.**
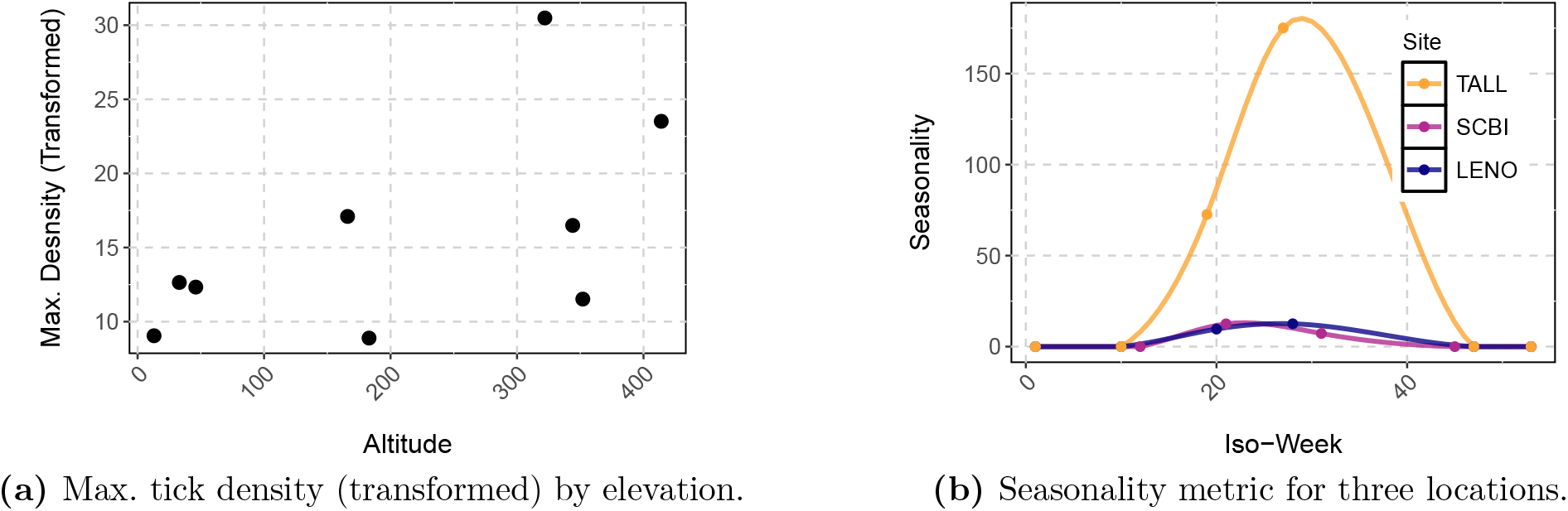
(a) and (b) represent location-specific predictors used in the GP model.

Lastly, we build a seasonality metric 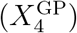, which uses foliage data obtained at each location (Site Metadata, 2025). We use the days-of-year (DOYs) when greenness increases, peaks, decreases, and reaches its minimum. These DOYs are converted to iso-weeks, 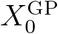, and are paired with corresponding tick densities *Y* . Two more data pairs for week 1 and 53, with density set to zero are added for continuity across years. This results in six pairs, 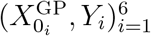, for each location. Then, we fit a cubic spline through these points as shown in Fig. 6b for the subset of locations shown previously in Fig. 2 (see Appendix S1: Section B for one detailed location). Lastly, we use the spline to make predictions *Y*_*i*_ for all 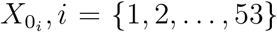 and map it the corresponding week at that location, creating 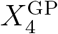.

This predictor follows the same trend as foliage, i.e., it increases in the summer and decreases in the winter. Each kernel (spline fit) of the seasonality metric is periodic but has a different height which allows us to incorporate some information about the tick density of one location relative to another. For example, when the magnitudes of the seasonality metric are closer together (i.e., iso-weeks 1–10, 45 – 53), the correlation in the responses between these locations is higher. When the magnitudes vary (at iso-week 20–30), the responses will be less similar. Another speciality of this predictor (unlike the sinusoid) is that the peak and the rate of change of the seasonality kernel across iso-weeks is allowed to vary from one location to another.

Using these predictors, we fit two GP models, (i) using data only obtained at one location namely *GP(L)* model through time dependent predictors (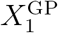 and 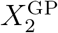 ) and (ii) using data obtained at all locations, *GP(A)* using both time and location predictors 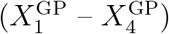. In order to fit the GP, the input space is scaled so all predictors lie within the interval [0,1]. We used an anisotropic GP (Eqs. 2–4) and fit the models using `hetGP` (Binois and Gramacy, 2021) package in R (CRAN).

In Fig. 7, we show fits and predictions of both *GP(L)* and *GP(A)* fits at BLAN from 2015 to 2022. Observe that *GP(L)*, as trained only on data obtained at BLAN, cannot adequately learn a trend and regresses towards the mean, possibly due to inadequate data. By contrast, *GP (A)* model “borrows” information from other locations and learns the seasonal pattern, i.e., higher tick abundance during summer and little to no activity in winter. Note that the upper bound of the *GP(L)* model is much smaller than *GP(A)* model. This is because the *GP(A)* is the average of the noise across all locations resulting in overestimation of noise at some locations such as BLAN.

**Figure 7.**
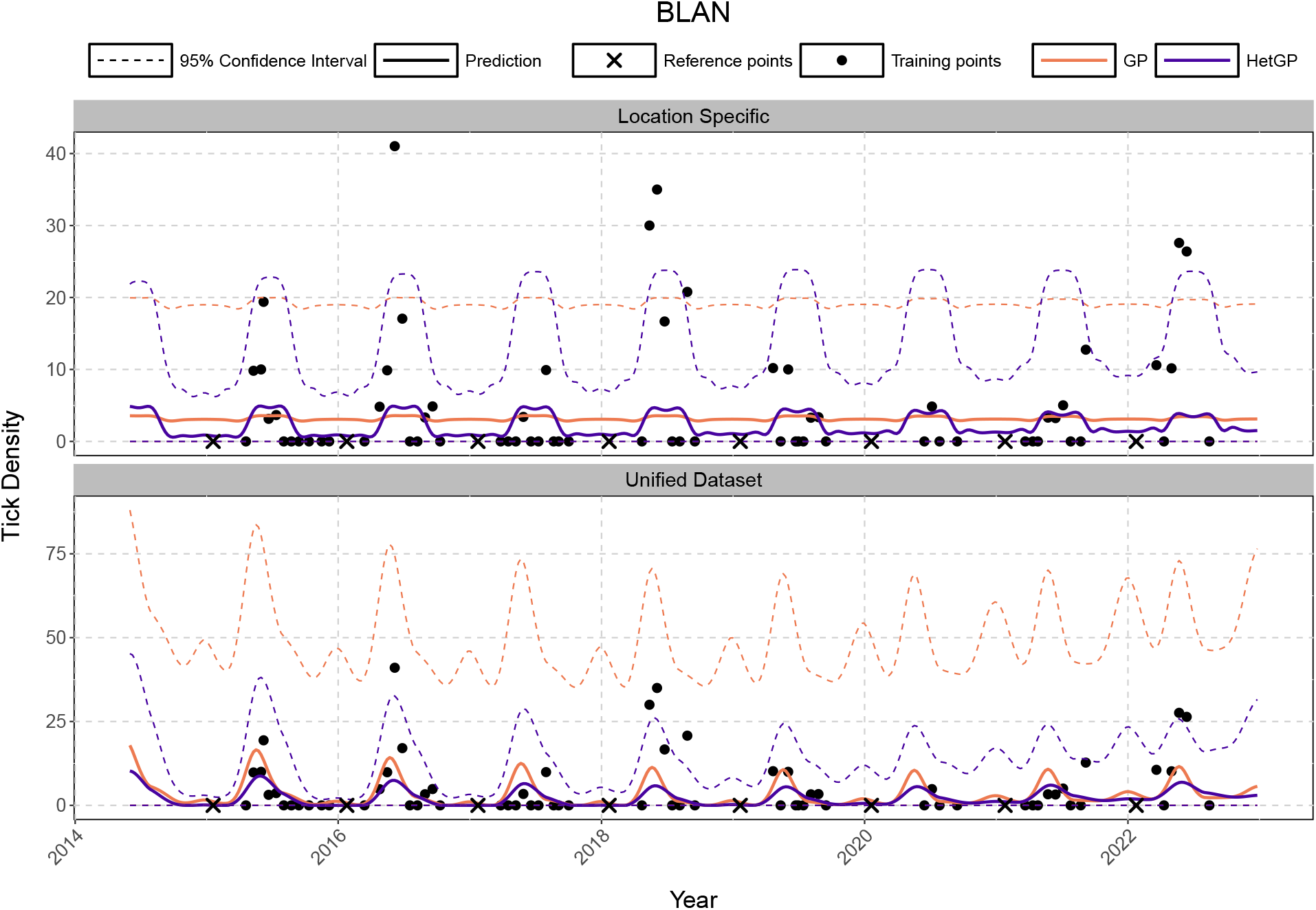
GP (orange) and HetGP (blue) fits at BLAN (solid = mean, dashed = 90% prediction intervals) on the original (untrasformed) scale. Black dots show training data; (*top)* models fit using location specific data only – *GP(L)* and *HetGP(L)*; (*bottom) GP(A)* and *HetGP(A)* trained on data from all locations.

### 3.3 Heteroskedastic Gaussian processes

In the previous sections, we have observed that the response varies across different locations (see Fig. 2). All the models discussed so far assume constant variance, which can lead to overinflation of noise at some locations with smaller densities (see Fig. 7) and vice-versa. To avoid compromising on UQ, we relax the assumption of constant noise across the input space and allow some flexibility in noise level estimation via a heteroskedastic Gaussian processes (HetGP; Binois et al., 2018). HetGPs model the mean process similar to a GP but additionally infer a latent noise process. Specifically, the model assumes that the noise changes across the input space. Mathematically, this corresponds to replacing *g*𝕀 → Λ_*n*_ in Eq. 2 so 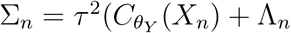 where, Λ_*n*_ = Diag(*λ*_1_, …, *λ*_*n*_) and

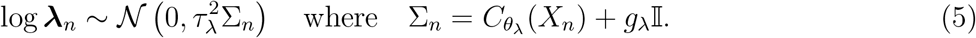

That is, the log-noise process follows a homoskedastic GP with the same kernel structure as in Eq. (2) but a different set of hyper-parameters, 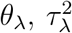, and *g*_*λ*_. The predictions follow Eqs. (3–4), with the new Σ_*n*_ matrix. To fit the hetGP model, we use the same predictors as the GP scaled to unit-cube and use the `hetGP` library on CRAN. We refer to this model as *HetGP(A)*.

Referring back to Fig. 7, observe that *HetGP(L)* captures a mean trend, but is unable to capture varying noise across the input space. At other locations, we also see *HetGP(L)* predictions regressing towards the mean (Appendix S1: Section C). By contrast, *HetGP(A)* trained on all available data provides better UQ. The bounds are tighter in winter where the model confidently predicts no tick activity but the UQ increases during summer months when there is higher noise. We provide a full comparison of percentage coverage along with interval widths in Section 4 showing that the *HetGP(A)* outperforms all models with appropriate bounds.

Since the *GP(L)* and *HetGP(L)* models are hit-or-miss, we focus on *GP(A)* v/s *HetGP(A)* to explain noise process estimation. The predictive variance is a combination of the process uncertainty (variability in the response) and noise (unexplained variation), which we suspect varies from one location to another and across seasons. GPs average across the noise in the input space using one noise level, *g*. This often results in over/under estimation (e.g., BLAN). HetGPs provide different noise level estimates across every week at every location. This results in adaptive prediction intervals that effectively capture input dependent noise and improve uncertainty estimation.

We use Fig. 8 to demonstrate the capabilities of *HetGP(A)*. We have two locations where the *GP(A)* and *HetGP(A)* provide the same coverage. Notice in Fig. 8a at SERC that noise estimated by the *GP(A)* model is consistently higher for the same coverage. At KONZ (Fig. 8c), we observe that *HetGP(A)* captures both seasonal and site-specific variation. For example, at KONZ during high-variability summer weeks (*n*^*′*^ ∈ { 18, …, 37 }), the model estimates higher noise (***λ****n*^*′*^ *> g*), leading to wider intervals whereas, in winter, noise is lower (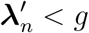 for *n*^*′*^ ∉ {14, …, 26, 29, …, 39 }), resulting in tighter bounds (Fig. 8d). This pattern is different than what is seen at SERC even though the locations are fit together with the same model. This adaptability enables *HetGP(A)* to provide better coverage and more reliable forecasts.

**Figure 8.**
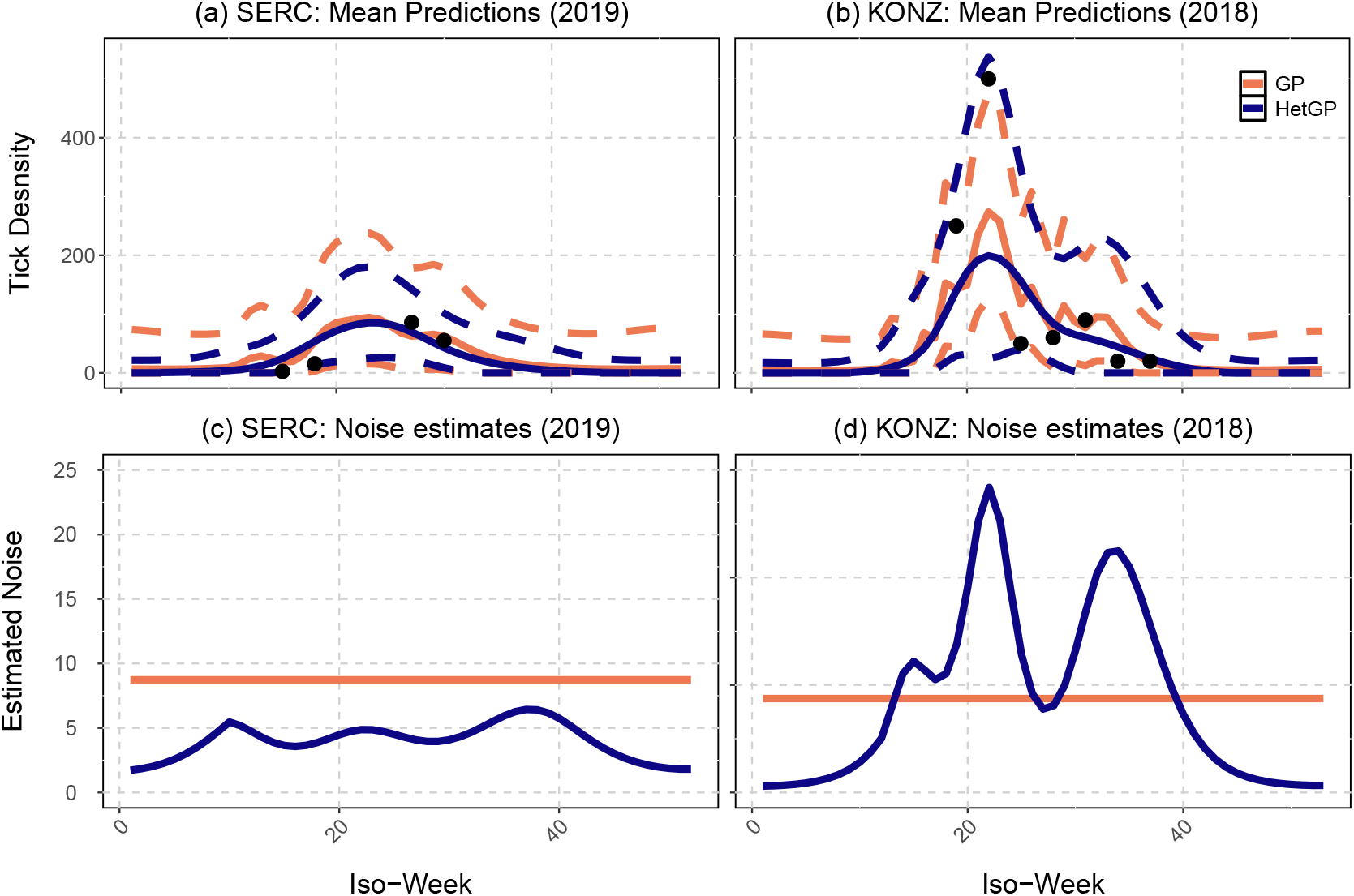
Predicted tick densities (solid = mean, dashed = intervals) and estimated noise levels for two locations obtained from regular GP (orange) and HetGP (blue) model.

## 4 Comparative Prediction Performance

In this section, we evaluate the performance of several models using metrics discussed in 2.2.2. We consider models such as LR-Temp, BASS, GP, and HetGP. We use two versions of all models: (1) Models trained using data at specific location (notated by L) using time dependent predictors such as week and periodicity and (2) models trained using all the available data which also use location specific predictors. For the *LR-Temp(A)* model, we use a categorical input to distinguish between locations. Lastly, we include *LR-Time(L)* model as our baseline.

We evaluate within-sample fits using data before 2023 by analyzing Q-Q plots of standardized residuals (Fig. 9). Residuals are computed using training data and model predictions, then standardized by dividing by the predicted standard error. The Q-Q plot compares predicted vs. observed tick densities, and closeness to the Q-Q line indicates better fit and UQ. GP models predict well at lower densities and stray away from the line at higher densities, slightly under-estimating tick abundance. All other models under predict at lower tick densities, which is arguably worse as detecting an absence of ticks when there are really ticks could lead to lack of preventive measures and increase in tick-borne diseases.

**Figure 9.**
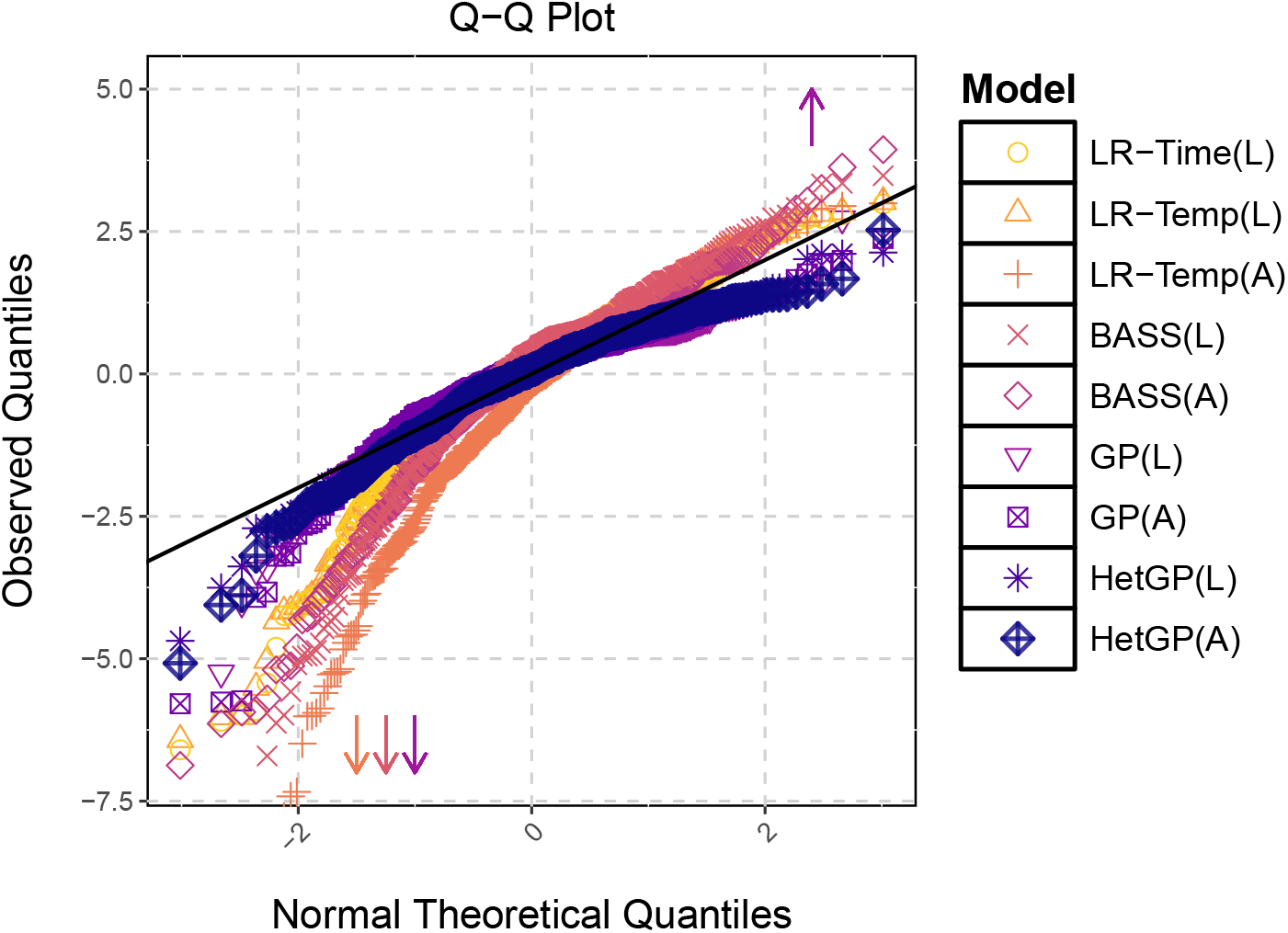
Q-Q Plot showing the standardized residuals of all models for all points in the training data

**Table 1.** Coverage (%) and interval widths for both in-sample and out-of-sample data for 90% prediction intervals.

| Model | In-sample |  |  | Out-of-sample |  |  |
| --- | --- | --- | --- | --- | --- | --- |
|  | Coverage (%) | Median Width | Mean Width | Coverage (%) | Median Width | Mean Width |
| LR-Time(L) | 95.49 | 10.90 | 15.18 | 87.90 | 13.15 | 17.22 |
| LR-Temp(L) | 95.49 | 11.21 | 11.77 | 87.90 | 11.49 | 11.92 |
| LR-Temp(A) | 91.93 | 14.02 | 14.09 | 89.14 | 14.06 | 14.11 |
| BASS(L) | 91.47 | 11.43 | 12.17 | 84.81 | 11.95 | 12.75 |
| BASS(A) | 92.17 | 10.95 | 10.95 | 87.90 | 10.97 | 10.98 |
| GP(L) | 95.04 | 10.21 | 10.49 | 92.59 | 12.66 | 12.79 |
| GP(A) | 94.24 | 10.33 | 10.49 | 89.07 | 11.37 | 11.56 |
| HetGP(L) | 91.55 | 9.76 | 9.76 | 86.05 | 13.09 | 13.25 |
| HetGP(A) | 94.15 | 8.04 | 8.34 | 89.44 | 10.27 | 10.76 |

We assess sample coverage and interval widths to evaluate whether prediction intervals provide adequate UQ. These metrics were calculated on the transformed predictive bounds to capture the full width of the intervals (back transforming results in lower PIs bounded at zero). The coverage of all models along with the mean and median width of the interval across all locations, for all observations is shown in Tab. 1. The in-sample coverage is typically between 90 − 95% for a 90% prediction interval for all models whereas, the out-of-sample coverage is between 85 − 90% with only the *GP(A)* providing *>* 90% coverage. Notice that *HetGP(A)* is closest to the nominal level for out-of-sample predictions and has smaller widths than all other models for in- and out-of sample data.

Lastly, we also evaluate in- and out-of-sample RMSE and CRPS. The RMSE and CRPS for the *HetGP(A)* model is generally lower at all locations compared to all other models. One situation where the *HetGP(A)* does not convincingly outperform other models is at UKFS in the testing (out-of-sample) set. Here, the magnitude for the testing data responses were much larger than previously recorded at the location. Additionally, the site is very sparsely sampled with only 25 total observations (Fig. 1). The *HetGP(A)* model simply learns patterns in historical data and forecasts these into the future with an assumption of stationarity. The results can sometimes be poor when there is a drastic change in the range of the response not present in historical training data. Also, recall Fig. 7 where *GP(L)* model clearly does not learn a pattern and simply provides a linear mean prediction. We notice similar behavior in the *BASS(L)* model at other locations. Judging purely on basis of metrics, we see all GP, HetGP, and BASS seem to perform similarly at BLAN, even despite the addition of reference points. Yet only *HetGP (A)* captures both the mean and noise process as evident in the Fig. 7.

## 5 Discussion

We proposed using Gaussian processes to forecast tick populations for one future season (here one year) and have shown their capability in settings with sparse data. Our framework leveraged a core property of GPs: reliance on relative distances between predictor/input values rather than their position in the input space. Additionally, we show that modeling noise separately via HetGPs provides better UQ and coverage. Ultimately we observed that the HetGP models outperformed all the other competitors.

Despite these advantages, GP models retain some of the challenges typical of more common time-series approaches. For example, GP models are best suited for short-term forecasting (e.g., one season/year or less) as these models typically assume stationarity which may not hold for longer horizons. GPs also do not include mechanistic components that describe the dynamics of the population being modeled. Unlike differential equation based models which study rate of change of population explained by factors such as demographics, temperature, or humidity, GP models do not posit either mechanisms or causality. Thus, they cannot be used to explore possible changes in populations across time due to climactic shifts or to implement control measures.

Understanding the noise process was essential to obtain better predictions. However, the current HetGP implementation lacks flexibility in specifying how noise varies across input dimensions. In ecological datasets, noise levels often fluctuate due to seasonal and geographical factors. In this paper, the HetGP model fits an individual noise level at every point in time across all locations. Although this worked well in our case, it may lead to overfitting in some scenarios. Future work could explore alternative implementations where the noise process only changes in some dimensions of the input space.

The goal of our case study was to develop a short-term forecasting model for a sparse dataset such as ticks. These forecasts provide insights into tick patterns and enable health departments and policy makers to take action and ensure disease prevention (Kessler et al., 2019; Goddard and Varela-Stokes, 2009). We can protect wildlife, livestock and pet infections (Sykes, 2023; Springer et al., 2015). However, we see broader applications using the GP framework. Using a similar recipe to ours, ecologists can use this model on other irregularly sampled datasets with large gaps such as mosquitoes, endangered species, etc.

## Appendix A Performance Metrics

1. **Residual Mean Square Error (RMSE)** RMSE provides a measure of prediction error. Let *y*_*i*_ be the response observed at input location *x*_*i*_ and *µ*_*i*_ denote the prediction at that location. Let *n* be the sample size. Then the RMSE can be given as follows:

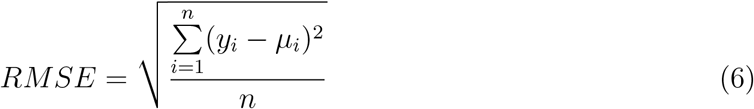
2. **Continuous Rank Probability Score (CRPS)** CRPS provides a measure of uncertainty quantification. Let *y*_*i*_ be the response with posterior predicted mean *µ*_*i*_ and standard deviation *σ*_*i*_. The CRPS is defined as follows:

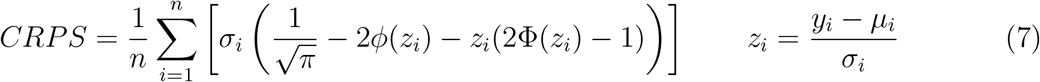

## B Spline Fitting for Seasonality Metric

Here, we provide a detailed explanation of how we fit a spline to obtain the seasonality metric for one location. (TALL)

1. Obtain points for one location to fit the spline. This is based on the foliage information provided by NEON.
2. Use 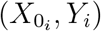 pairs and fit a spline using R cubic.spline = spline(X,Y, n=53, method = “natural”) Here n = 53 indicates we want 53 points which will be X = 1, 2 …53 and corresponding Y.
3. Obtain *Y* for all *X* = 1, 2, ….53 from the fitted spline. If any Y is negative, we bound it at 0. This gives us a positive seasonality metric.

**Table 2.** Fitting a spline for Seasonality Metric; Section 3.2 (Fig. 6).

| Foliage Trend | DOY | $X_0$ | Y | $(X_{0_i}, Y_i)$ |
| --- | --- | --- | --- | --- |
| Continuity |  | 1 | 0 | (1, 0) |
| Starts increasing | 75 | 10 | 20 | (10, 20) |
| Reaches Maximum | 135 | 19 | 174.85 | (19, 174.85) |
| Starts Decreasing | 195 | 27 | 55.08 | (27, 55.08) |
| Reaches Minimum | 330 | 47 | 0 | (47, 0) |
| Continuity |  | 53 | 0 | (53, 0) |

## C Location-wise Data Visualization and Model Fitting

In this section, we show the fits obtained from all the models: *LR-Time, LR-Temp, BASS, GP*, and *HetGP*, at each location. We include both the *(L)* and *(A)* methods trained using a subset and all of the data respectively. Observe that at locations which is very irregularly sampled (e.g., BLAN, LENO, etc. ), *HetGP(L)* and *BASS(L)* also suffer from mean regression (similar to the *GP(L)* model described in Fig. 7). Additionally, we observe that training *LR-Temp(A)* and *BASS(A)* using all the available data also behaves similar to the *GP(A)* averaging across the noise over all locations resulting in overly conservative bounds at some locations (e.g., BLAN) and very poor coverage at others (e.g., UKFS).

**Figure 10.**
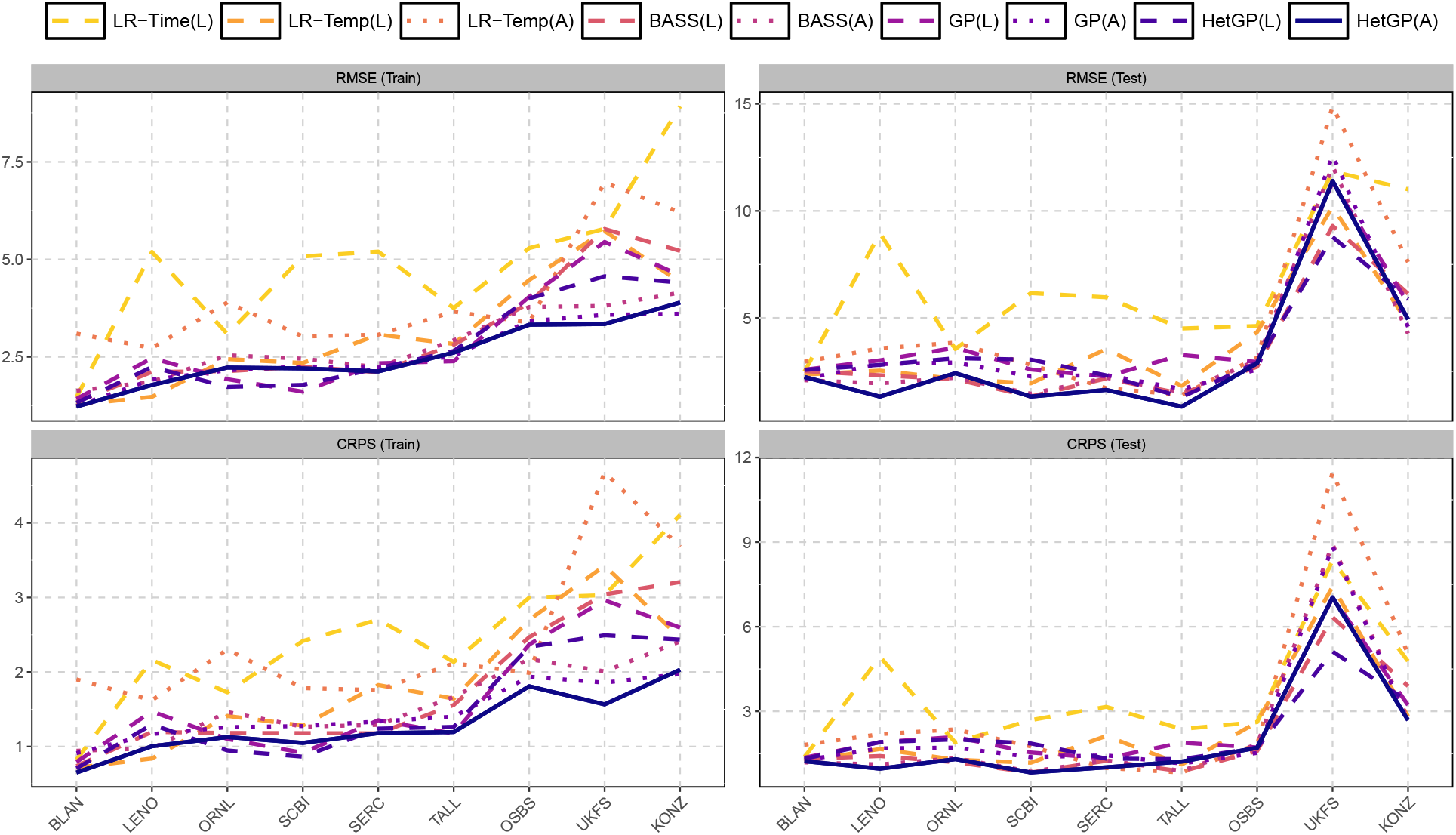
Train and test CRPS and RMSE. Models trained only using location-wise data are in *dashed* and on all locations in *dotted*. Our approach: *HetGP* model is shown in *solid* purple line.

**Figure 11.**
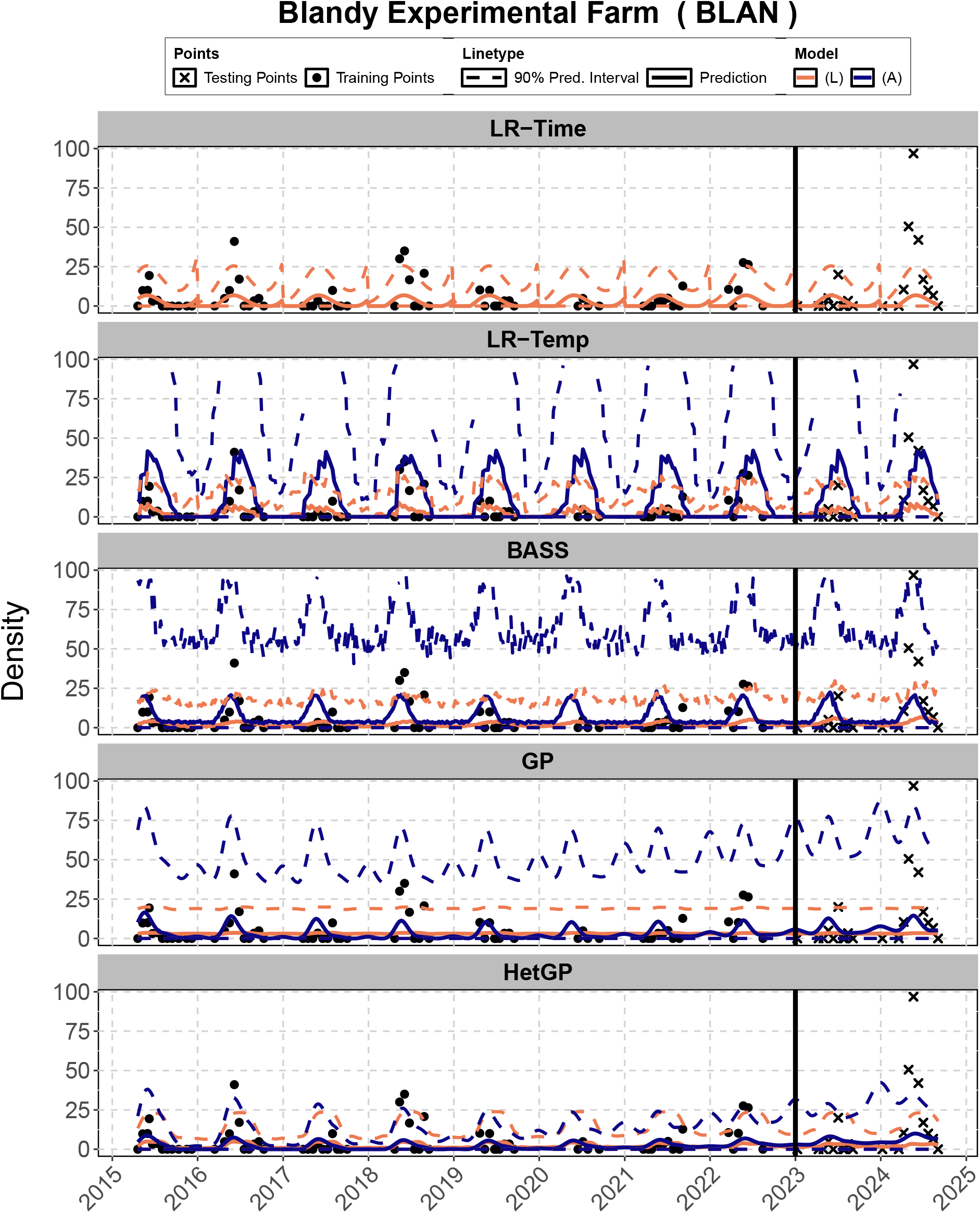
*LR-Time, LR-Temp, BASS, GP*, and *HetGP* – (L) and (A) models at BLAN along with 90% prediction intervals on training and testing dataset

**Figure 12.**
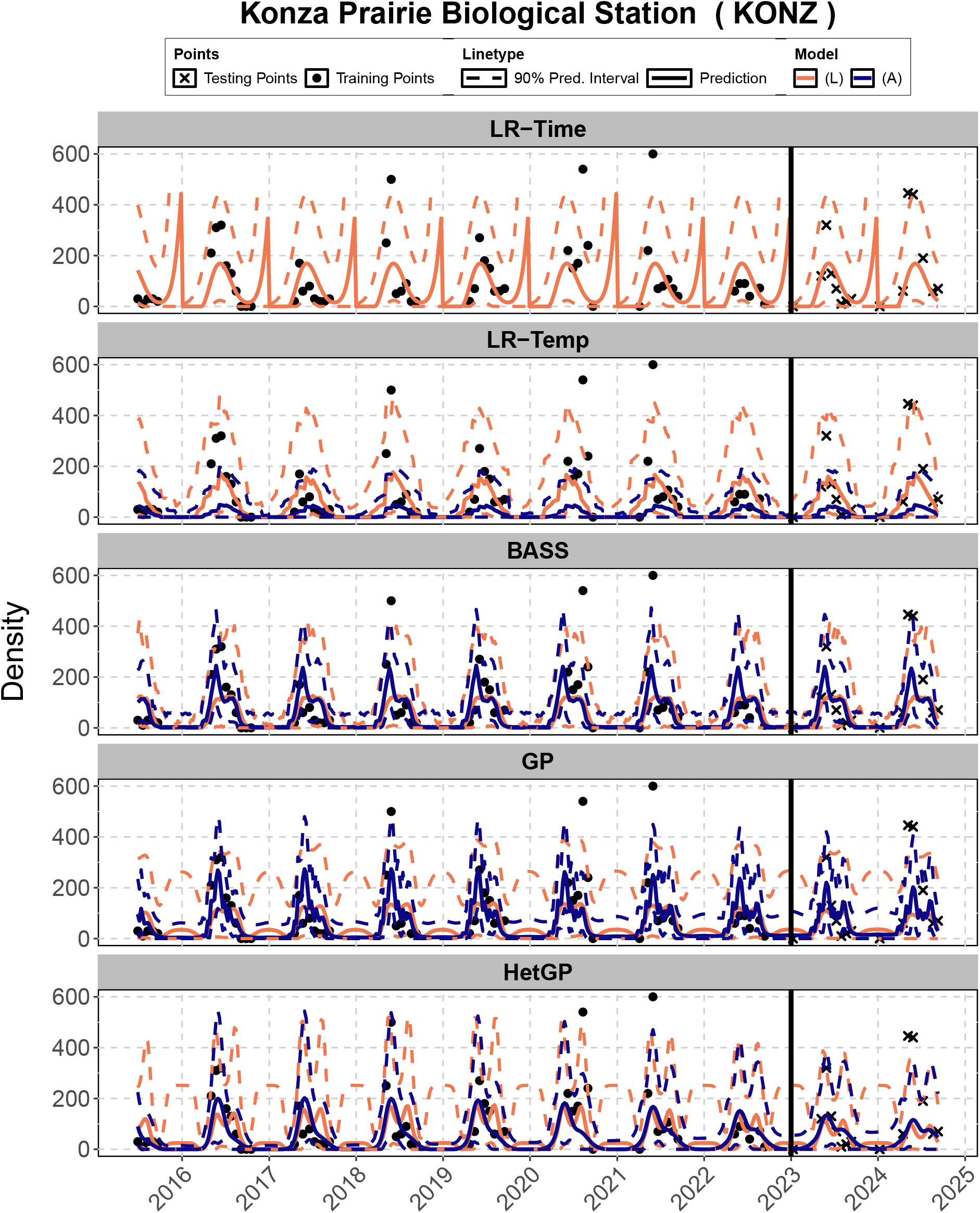
*LR-Time, LR-Temp, BASS, GP*, and *HetGP* – (L) and (A) models at KONZ along with 90% prediction intervals on training and testing dataset

**Figure 13.**
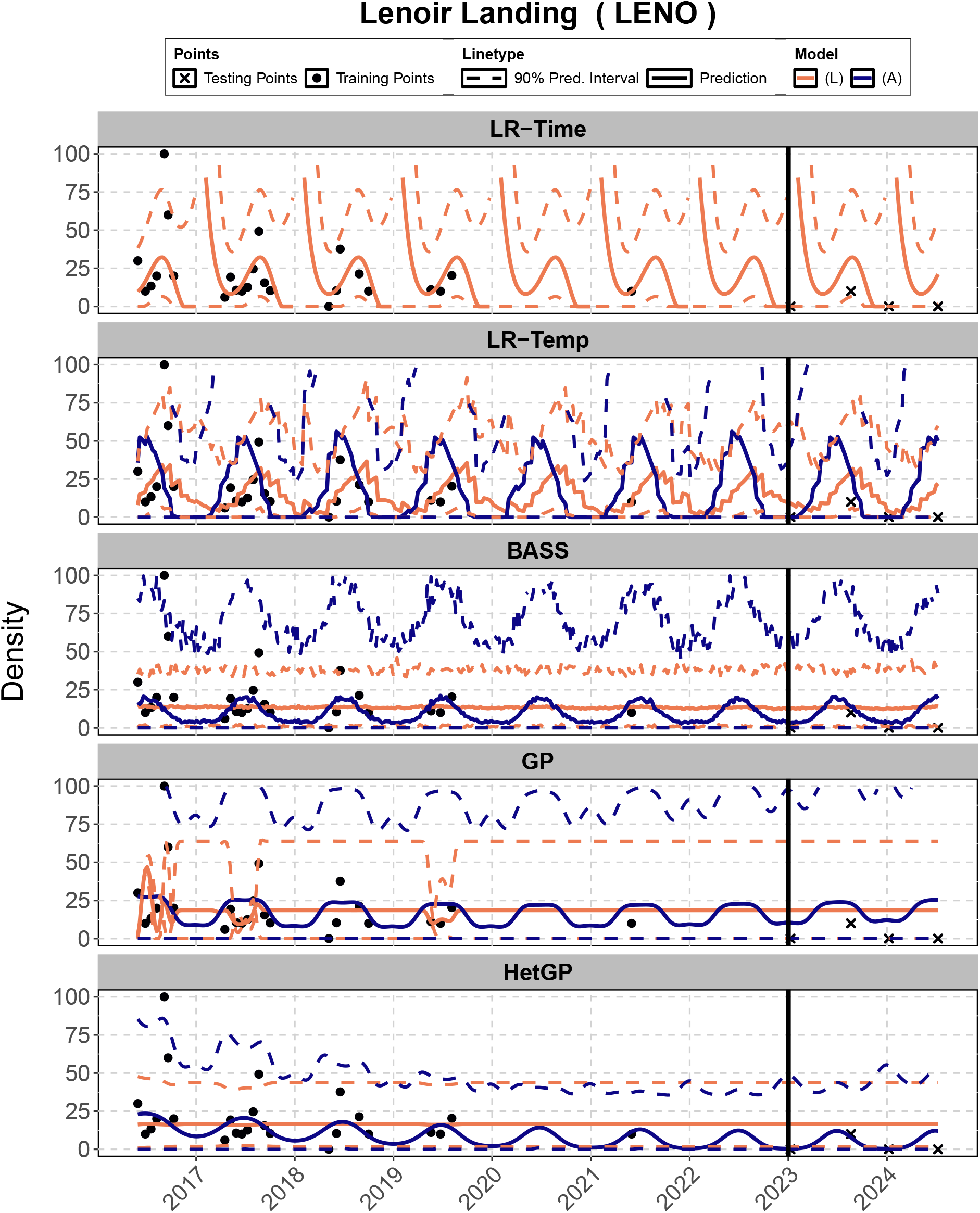
*LR-Time, LR-Temp, BASS, GP*, and *HetGP* – (L) and (A) models at LENO along with 90% prediction intervals on training and testing dataset

**Figure 14.**
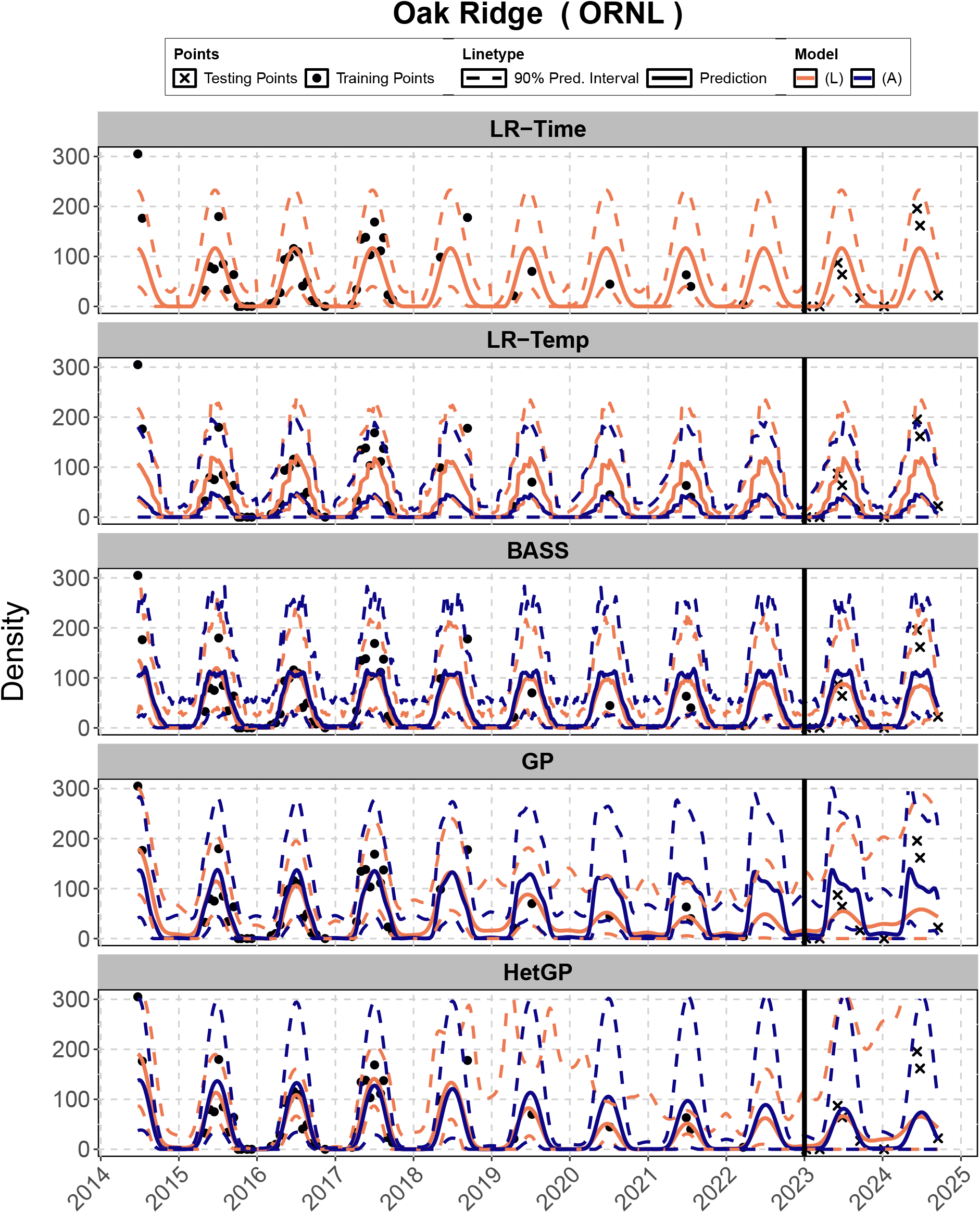
*LR-Time, LR-Temp, BASS, GP*, and *HetGP* – (L) and (A) models at ORNL along with 90% prediction intervals on training and testing dataset

**Figure 15.**
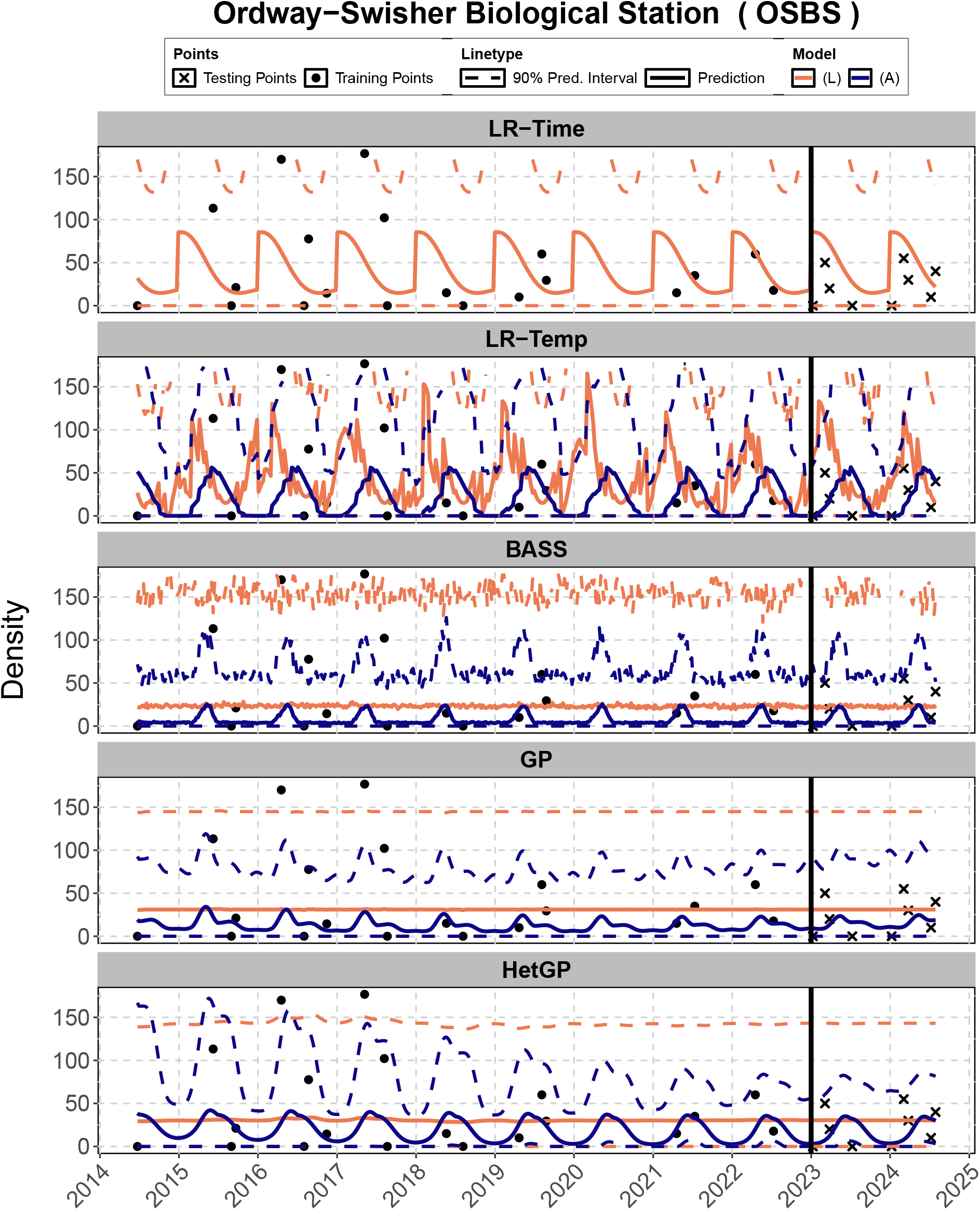
*LR-Time, LR-Temp, BASS, GP*, and *HetGP* – (L) and (A) models at OSBS along with 90% prediction intervals on training and testing dataset

**Figure 16.**
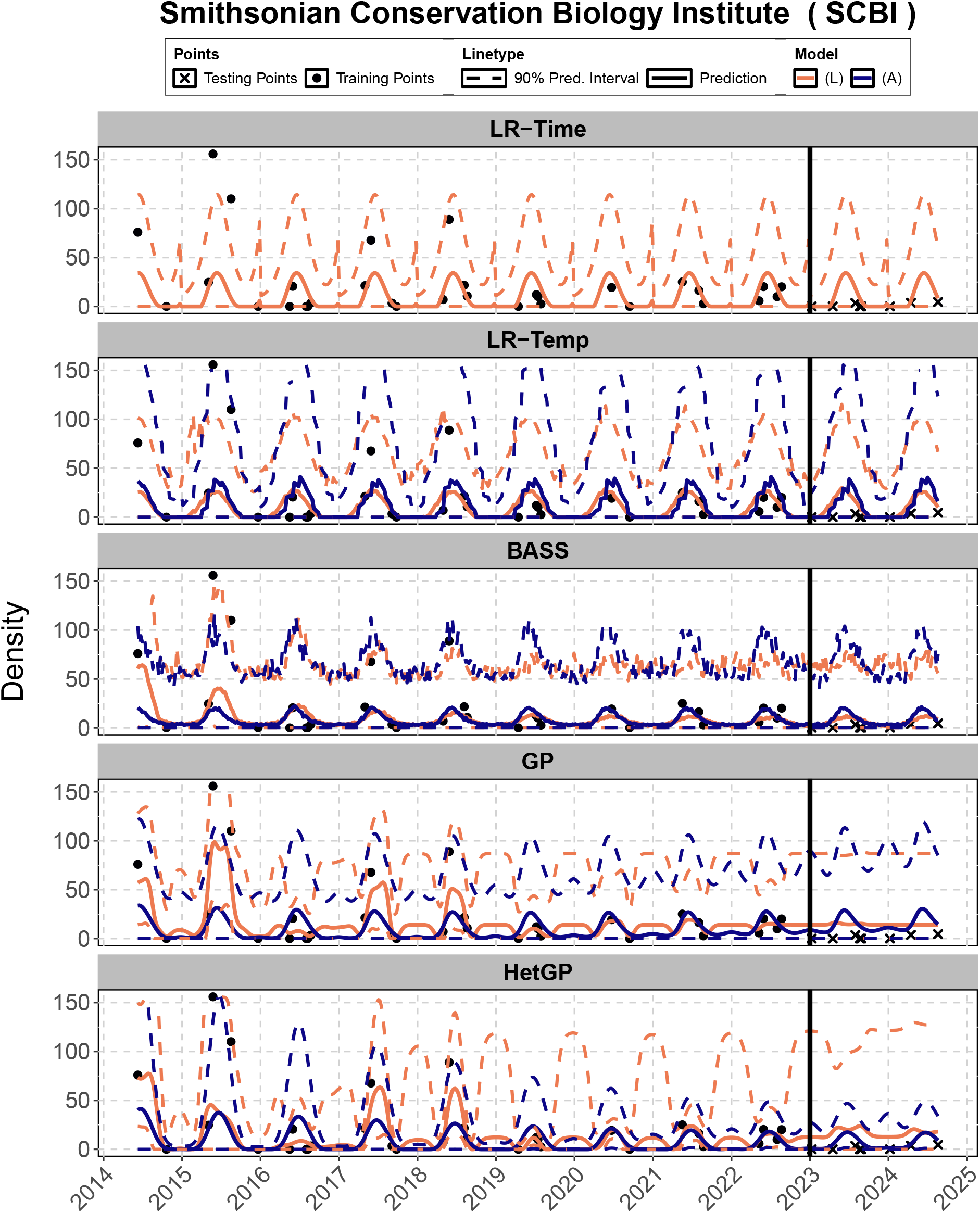
*LR-Time, LR-Temp, BASS, GP*, and *HetGP* – (L) and (A) models at SCBI along with 90% prediction intervals on training and testing dataset

**Figure 17.**
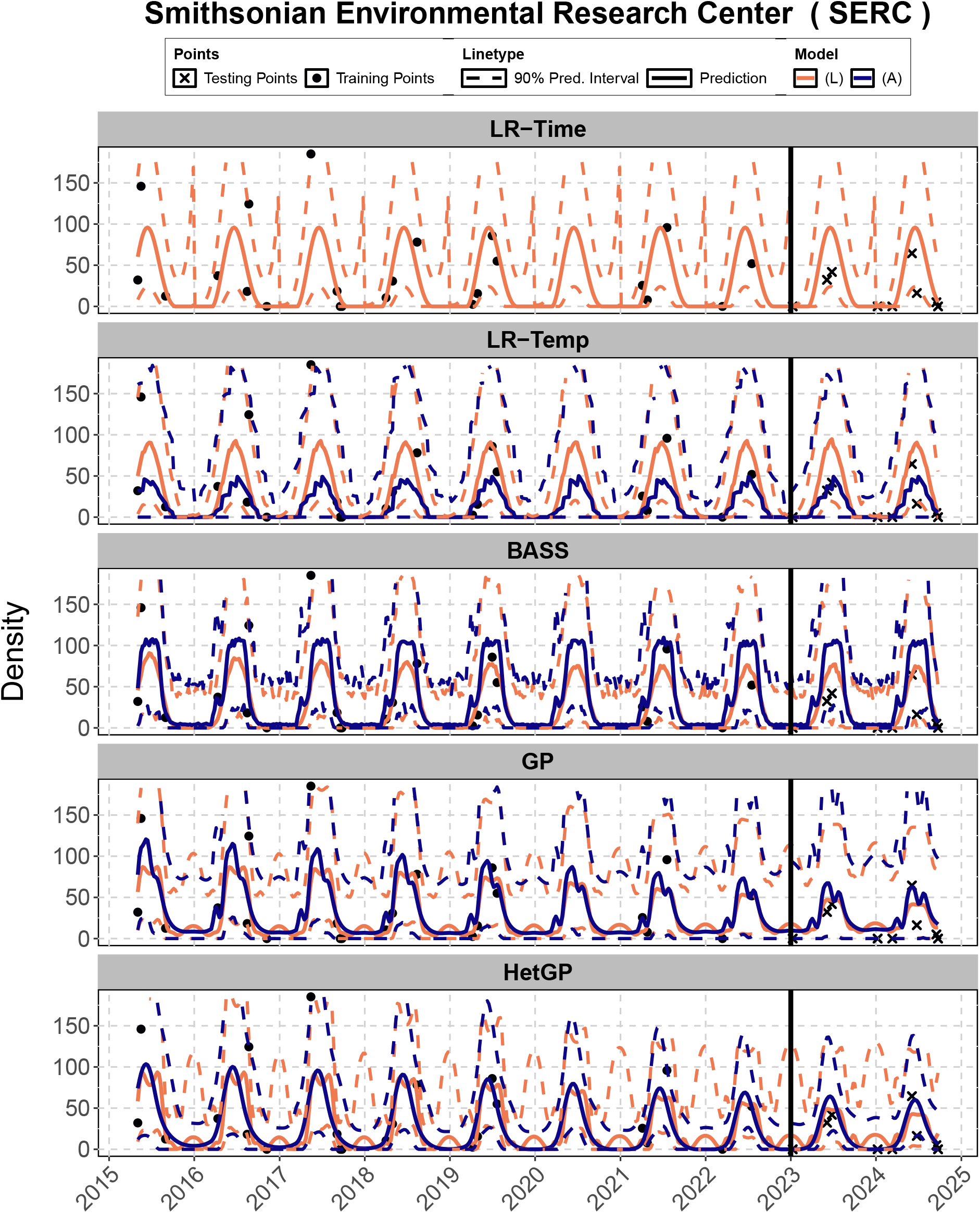
*LR-Time, LR-Temp, BASS, GP*, and *HetGP* – (L) and (A) models at SERC along with 90% prediction intervals on training and testing dataset

**Figure 18.**
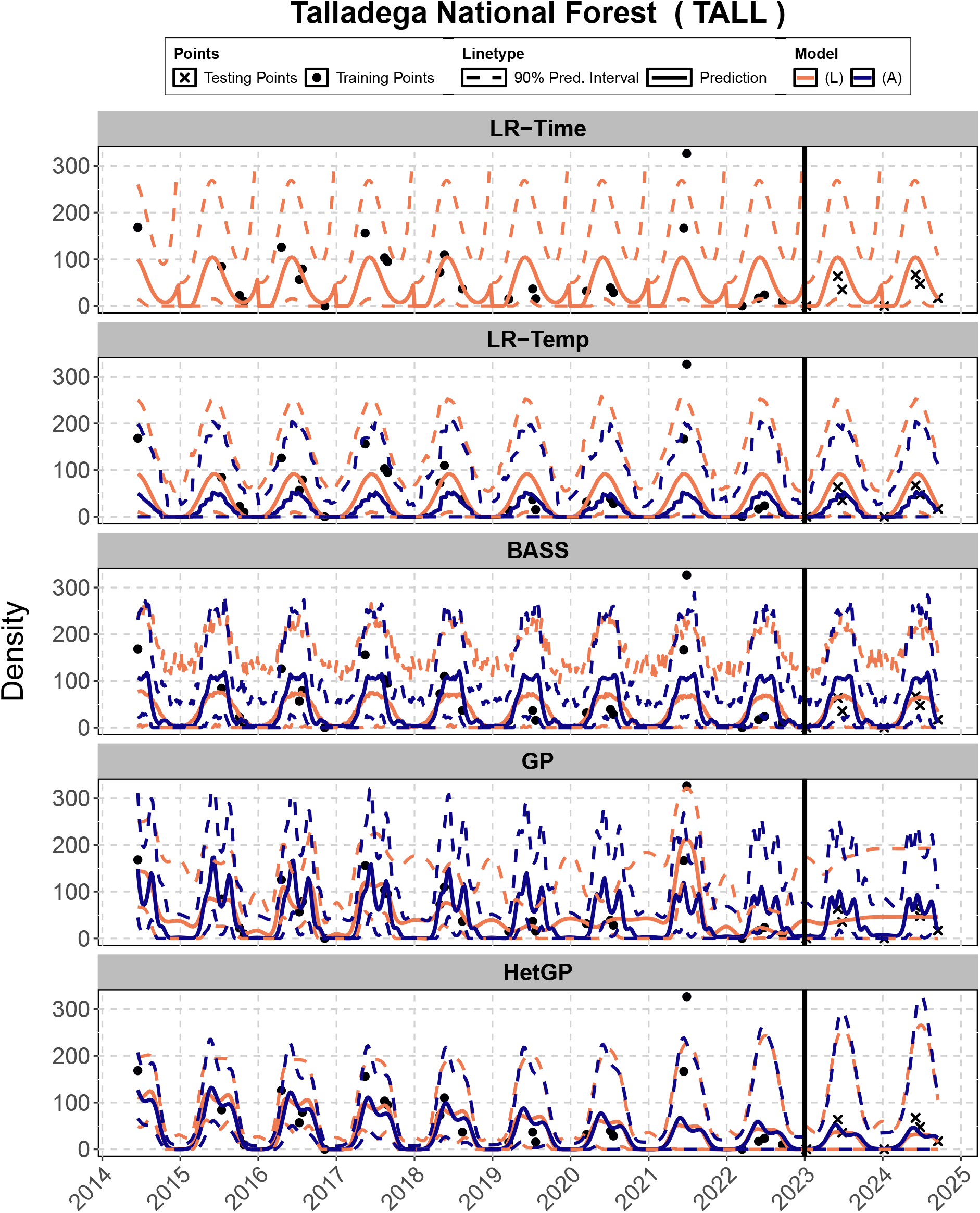
*LR-Time, LR-Temp, BASS, GP*, and *HetGP* – (L) and (A) models at TALL along with 90% prediction intervals on training and testing dataset

**Figure 19.**
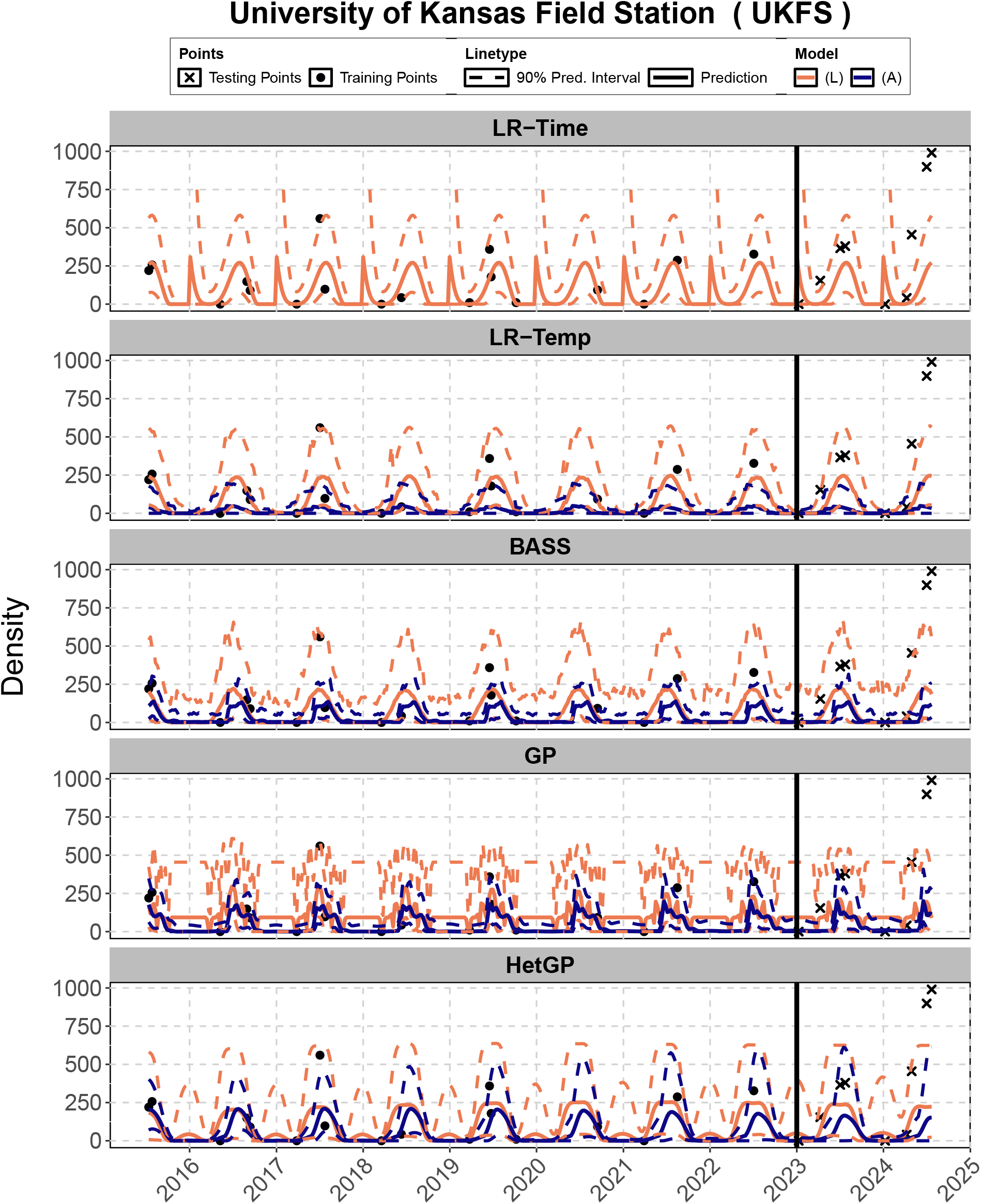
*LR-Time, LR-Temp, BASS, GP*, and *HetGP* – (L) and (A) models at UKFS along with 90% prediction intervals on training and testing dataset

## Footnotes

1 See https://www.ncei.noaa.gov/products/weather-climate-models/global-ensemble-forecast.

